# Connectome predictive modeling of trait mindfulness

**DOI:** 10.1101/2024.07.09.602725

**Authors:** Isaac N. Treves, Aaron Kucyi, Madelynn Park, Tammi R.A. Kral, Simon B. Goldberg, Richard J. Davidson, Melissa Rosenkranz, Susan Whitfield-Gabrieli, John D.E. Gabrieli

**Author notes:** **Corresponding author:** Isaac Treves;, Building 46-4037, Cambridge, MA, 02139. Conflict of interest statement: The authors declare no competing interests. Ethics statement: Procedures for the Wisconsin study were approved by the Healthy Sciences Institutional Review Board of the University of Wisconsin-Madison. Procedures for the Leipzig study were approved by the ethics committee at the medical faculty of the University of Leipzig (097/15-ff). The Stanford study was approved by the Stanford University Institutional Review Board.

## Abstract

**Introduction:** Trait mindfulness refers to one’s disposition or tendency to pay attention to their experiences in the present moment, in a non-judgmental and accepting way. Trait mindfulness has been robustly associated with positive mental health outcomes, but its neural underpinnings are poorly understood. Prior resting-state fMRI studies have associated trait mindfulness with within- and between-network connectivity of the default-mode (DMN), fronto-parietal (FPN), and salience networks. However, it is unclear how generalizable the findings are, how they relate to different components of trait mindfulness, and how other networks and brain areas may be involved.

**Methods:** To address these gaps, we conducted the largest resting-state fMRI study of trait mindfulness to-date, consisting of a pre-registered connectome predictive modeling analysis in 367 adults across three samples collected at different sites.

**Results:** In the model-training dataset, we did not find connections that predicted overall trait mindfulness, but we identified neural models of two mindfulness subscales, *Acting with Awareness* and *Non-judging*. Models included both positive networks (sets of pairwise connections that positively predicted mindfulness with increasing connectivity) and negative networks, which showed the inverse relationship. The *Acting with Awareness* and *Non-judging* positive network models showed distinct network representations involving FPN and DMN, respectively. The negative network models, which overlapped significantly across subscales, involved connections across the whole brain with prominent involvement of somatomotor, visual and DMN networks. Only the negative networks generalized to predict subscale scores out-of-sample, and not across both test datasets. Predictions from both models were also negatively correlated with predictions from a well-established mind-wandering connectome model.

**Conclusions:** We present preliminary neural evidence for a generalizable connectivity models of trait mindfulness based on specific affective and cognitive facets. However, the incomplete generalization of the models across all sites and scanners, limited stability of the models, as well as the substantial overlap between the models, underscores the difficulty of finding robust brain markers of mindfulness facets.

---

Mindfulness, often defined as the act of paying attention to the present moment without judgement (Bishop et al., 2004), is a construct that has been researched intensely in recent years. Researchers have sought to operationalize mindfulness as a trait, known as *trait mindfulness*, which refers to one’s disposition or tendency to pay attention to experiences in a mindful way. Trait mindfulness is often measured by self-report scales, such as the Five Facet Mindfulness Questionnaire (FFMQ) (R. A. Baer et al., 2004) and the Mindful Attention Awareness Scale (MAAS) (Brown & Ryan, 2003). Greater trait mindfulness has been frequently associated with a range of positive mental health outcomes, including more positive affect, improved self-compassion, greater openness to experience, and better quality of life (Allen et al., 2021; Amundsen et al., 2020; Chu & Mak, 2020; Kong et al., 2014; Schutte & Malouff, 2011), and negatively associated with outcomes like negative affect, stress, and anxiety (Carpenter et al., 2019; Coffey & Hartman, 2008; de Bruin et al., 2014; Greco et al., 2011; Tomlinson et al., 2018; Treves et al., 2023). Given the importance of trait mindfulness as a predictor of mental health, there is a clear need to understand its neural underpinnings. Neuroimaging of trait mindfulness could help us understand mental health conditions (Zhuang et al., 2017), reveal pathways of action in mindfulness interventions (Goldberg et al., 2019), and provide possible targets for neuromodulation (Cain et al., 2024; Zhang et al., 2023). Motivated by these possibilities, this study investigated the functional neuroimaging basis of trait mindfulness.

Resting-state functional magnetic resonance imaging (fMRI) data, measured when participants lie awake in the fMRI scanner in a task-free state, may provide brain-based measures correlating with trait mindfulness. A resting state does not explicitly engage cognitive or emotional processes. Instead, it is used to study correlated self-generated brain signals, or functional connectivity, while a participant is at rest. These correlations, which can be reliable given sufficient data (Finn et al., 2015; Laumann et al., 2015; Noble et al., 2017), are thought to reflect stable aspects of individual functional brain organization (Shen et al., 2017; Smith et al., 2013). In addition, resting state data are relatively easy to collect and can be compiled in large online databases to increase sample size (Biswal et al., 2010; Eickhoff et al., 2016; Poldrack & Gorgolewski, 2017).

Static functional connectivity (SFC) is measured by correlations between brain regions over the course of a resting-state fMRI scan, and several networks of correlated brain regions are plausibly related to variation in trait mindfulness. One network is the default-mode network (DMN), involved in internally-focused, self-referential processing, and consisting of brain areas such as the precuneus, posterior cingulate, and ventromedial prefrontal cortex (Raichle et al., 2001). Two other candidate networks are the salience network (SN), involved in stimulus-driven attention and including the insula and mid-cingulate (Seeley et al., 2007); and the frontoparietal network (FPN), involved in externally focused, goal-directed attention and consisting of lateral frontal and parietal areas (Dosenbach et al., 2008; Greicius et al., 2003; MacDonald et al., 2000).

Despite at least nine studies on resting-state SFC and trait mindfulness (Bilevicius et al., 2018; Doll et al., 2015; Harrison et al., 2019; Hunt et al., 2022; Kong et al., 2016; Li et al., 2022; Parkinson et al., 2019; Shaurya Prakash et al., 2013; Wang et al., 2014), there is no consistent relation between these networks and trait mindfulness. For example, Bilevicius et al. (2018) found that *decreased* connectivity of the SN and the cuneus (often considered part of the DMN) correlated with MAAS total scores, but Parkinson et al. (2019), found that *increased* connectivity of the SN and cuneus correlated with FFMQ total scores. Both studies were conducted with *n* ∼30 participants. Some studies found that trait mindfulness correlated with reduced within-DMN connectivity (Bilevicius et al., 2018; Doll et al., 2015; Harrison et al., 2019; Wang et al., 2014), but a subsequent larger sample study (*n∼*100) failed to replicate this finding (Hunt et al., 2022). Sources of variability between studies could stem from variable sample characteristics, small sample sizes, different methodologies (i.e., choice of seed regions), or a lack of test-retest reliability in the fMRI measures. A broader concern is that mindfulness may not be a unitary trait (Altgassen et al., 2023; Beloborodova & Brown, 2023) and, therefore, it is unlikely to involve unitary brain processes. It may be that mindfulness involves related but distinct subcomponents like attention and non-judgement (Bishop et al., 2004). Indeed, the Five Facet Mindfulness Questionnaire (FFMQ) was developed based on factor analyses of previous mindfulness questionnaires, resulting in statistically dissociable facets of *Acting with Awareness, Non-judging, Non-reactivity, Describing, and Observing* (Baer et al., 2006, 2008; although *Observing* may show limited validity, Gu et al., 2016). These individual components may relate to different patterns of resting-state brain connectivity.

Here we addressed these concerns in a multisite study of resting-state fMRI and trait mindfulness to date. The present sample (*n* = 367) constitutes the largest sample size of any laboratory-based neuroimaging study of trait mindfulness (for a systematic review, see Treves et al., in press). We used a data-driven, whole-brain approach called connectome predictive modeling (CPM). CPM tests pairwise connections across the whole brain (Shen et al., 2017), and can find positive and negative network models that predict individual differences. A key feature of CPM is prediction – whereas correlation may inflate the strength of an association, prediction of held-out data is more accurate (Gabrieli et al., 2015). CPM has proven predictive power for individual differences in IQ, creativity, sustained attention, mind-wandering, and other traits (Beaty et al., 2018; Finn et al., 2015; Kucyi et al., 2021; Rosenberg et al., 2015). In this study, we used CPM to investigate relationships between trait mindfulness, assessed with the FFMQ, and resting-state SFC in functional networks across the whole brain (including the DMN, FPN, and SN). We assessed whether the relationships generalized to independent samples, and we examined the relations of brain networks to the overall FFMQ score as well as the five FFMQ facets. We had no *a priori* hypotheses given the inconsistencies in the previous literature.

## Methods

We preregistered this study before analysis at https://osf.io/dtk9a/. All deviations are reported in the **Supplement.**

## Training Dataset: Wisconsin

We obtained imaging and phenotypic data from the University of Wisconsin-Madison meditation study (NCT02157766). The sample consisted of 206 meditation-naïve participants (age *M* = 30.9, *SD* = 13.1 years, 85 male) who completed an eyes-open resting-state scan and the Five Facet Mindfulness Questionnaire (FFMQ) (Baer et al., 2006). Of those 206 participants, 71 had asthma, and the full sample was retained. One of the aims of the original trial was to evaluate the relationship between psychological factors and asthma, but this aim was not relevant to the present study. Asthma status was controlled for in our analyses by conducting partial correlations with an indicator variable, given evidence for relationships between chronic inflammatory conditions and functional connectivity (Aruldass et al., 2021; Labrenz et al., 2019) as well as mental health (Stanescu et al., 2019). No participants had psychiatric diagnoses.

Images were acquired on a GE MR750 3.0 Tesla MRI scanner with a 32-channel head coil. Anatomical scans consisted of a high-resolution 3D T1-weighted inversion recovery fast gradient echo image (450 ms inversion time; 256 × 256 in-plane resolution; 256 mm field of view (FOV); 192 × 1.0 mm axial slices). A 12 min functional resting-state scan run was acquired using a gradient echo echo-planar imaging (EPI) sequence (360 volumes; repetition time (TR)/echo time (TE)/Flip, 2000/20 ms/75°; 224 mm FOV; 64 × 64 matrix; 3.5 × 3.5 mm in-plane resolution; 44 interleaved sagittal slices; 3 mm slice thickness with 0.5 mm gap). The in-plane resolution was decreased after the first 21 participants from 3.5 × 3.5 to 2.33 × 3.5 mm to better address sinus-related artifacts, resulting in a matrix of 96 × 64. Resolution change was controlled for in subsequent analyses by partialling out an indicator variable.

### Test Dataset: Stanford Science of Behavior Change

We obtained imaging and phenotypic data from the Stanford Science of Behavior Change project (https://scienceofbehaviorchange.org/projects/poldrack-marsch/)(Bissett et al., 2023). The sample consisted of 82 meditation-naïve participants (age *M* = 23.6, *SD* = 4.9 years, 27 male) who completed an 8-min eyes-open resting state scan and the FFMQ (Baer et al., 2006). Of those 82 participants, 22 had diagnoses of anxiety, depression, or other clinical conditions, and all participants were retained, as results were no different when removing participants with diagnoses.

Participants were scanned in a GE Discovery MR750 3-Tesla system with a 32-channel Nova Medical head coil at the Stanford center for Cognitive and Neurobiological Imaging. The T1-weighted scan used a BRAVO sequence with the following parameters: duration (4 minutes and 50 seconds), TR (7.24ms), TE (2.784ms), flip angle (12°), slice number (186), and resolution (.9mm isometric voxels). The T2*-weighted, gradient-echo echo-planar imaging, scan parameters were as follows: duration (8 min), multiband acceleration factor (8), TR (0.68 s), TE (30 ms), flip angle (53°), echo spacing (0.57 ms), slice number (64), resolution (2.2 mm isotropic), phase encoding anterior to posterior.

### Test Dataset: Leipzig Mind-Brain-Body

We downloaded openly available imaging and phenotypic data from the functional connectome phenotyping dataset (Babayan et al., 2019), a component of the MPI-Leipzig Mind-Brain-Body project (Mendes et al., 2019). Procedures for this study were approved by the ethics committee at the medical faculty of the University of Leipzig (097/15-ff). The sample consisted of 79 meditation-naïve participants (modal age range 20-25, 45 male) who completed four eyes-open resting-state scans and completed the FFMQ (Baer et al., 2006), translated to German. No participants had psychiatric diagnoses.

Participants were scanned in a 3-Tesla Siemens Magnetom Verio system with a 32-channel head coil at the University of Leipzig. The T1-weighted, 3DMP2RAGE, scan parameters were as follows: duration (8.22 min), TR (5 s), TE (2.92 ms), flip angle 1/2 (4/5°), TI 1/2 (700/2500 ms), slice number (176), resolution (1.0 mm isotropic). The T2*-weighted, gradient-echo echo-planar imaging, scan parameters were as follows for each of the four runs: duration (15 min 30 s), multiband acceleration factor (4), TR (1.4 s), TE (39.4 ms), flip angle (69°), echo spacing (0.67 ms), slice number (64), resolution (2.3 mm isotropic). In the first and third runs, the phase encoding direction was anterior to posterior, whereas in the second and fourth runs, the phase encoding direction was posterior to anterior.

## Measures

The Five Facet Mindfulness Questionnaire consists of 39 questions, corresponding to five statistically separable subscales: *Acting with Awareness, Non-judging, Non-reactivity, Describing,* and *Observing* (Baer et al., 2006; Baer et al., 2008)*. Acting with Awareness (AA)* refers to attending to one’s actions in the present moment, e.g., “It seems I am running on automatic without much awareness of what I’m doing.” *Non-judging (NJ)* refers to not evaluating or judging one’s thoughts or feelings, e.g., “I criticize myself for having irrational or inappropriate emotions.” *Non-reactivity (NR)* is defined as allowing thoughts to come and go without being caught up in them, e.g., “I perceive my feelings and emotions without having to react to them.” *Describing (D)* refers to labeling experiences with words, e.g., “I am good at finding words to describe my feelings.” Finally, *Observing (OBS)* is defined as noticing or attending to internal experiences, e.g., “I pay attention to sensations, such as the wind in my hair or sun on my face.” Each question on the FFMQ is rated on a 5-point Likert scale, ranging from *“1 = Never or very rarely true” to “5 = Very often or always true”.* The *Acting with Awareness (AA), Non-judging (NJ), and Describing (D)* subscales include reverse-scored questions. The lowest possible total score is 39 and the highest possible score is 195, with higher scores representing higher levels of mindfulness. We used the total scores, the total scale without observing (Baer et al., 2022; Gu et al., 2016; Pang & Ruch, 2019), and the subscales. The FFMQ has demonstrated acceptable internal consistency across a range of samples (.72–.92, Baer et al., 2008). We assessed relationships between the subscales using Pearson’s correlations in the Wisconsin dataset. For comparisons of the total FFMQ scores between the datasets, we conducted simple unpaired, heteroskedastic t-tests.

## Procedure

Preprocessing was identical to Kucyi et al. (2021) and details are provided. We preprocessed each fMRI run individually using the same procedures across datasets, based on procedures implemented in the CONN toolbox (version 21a (https://www.nitrc.org/projects/conn) (Whitfield-Gabrieli & Nieto-Castanon, 2012) and SPM12 in Matlab R2019a (Mathworks Inc., Natick, MA). Preprocessing steps included deletion of the first four volumes, realignment and unwarping (Andersson et al., 2001), and identification of outlier frames (frame-wise displacement >0.9 mm or global BOLD signal change >5 SD) (Nieto-Castanon, 2020). Functional and anatomical data were normalized into standard MNI space and, in a unified step, segmented into gray matter, white matter (WM), and cerebrospinal fluid (CSF) (Ashburner & Friston, 2005). Smoothing of fMRI data consisted of spatial convolution with a Gaussian kernel of 6 mm full-width half-maximum (FWHM).

fMRI denoising involved linear regression of the following parameters from each voxel: (a) 5 noise components each from minimally-eroded WM and CSF, respectively, based on aCompCor procedures (Behzadi et al., 2007; Chai et al., 2012) (b) 12 motion parameters (3 translation, 3 rotation, and associated first-order derivatives); (c) all outlier frames identified within participants; and d) linear BOLD signal trend within session. After nuisance regression (Hallquist et al., 2013), data were bandpass filtered to 0.008–0.09 Hz. Denoising procedures have been shown to reduce the impact of head motion on functional connectivity (Muschelli et al., 2014), but excessive head motion may confound estimates (Power et al., 2015; Siegel et al., 2017). To avoid this possibility, we excluded participants with mean overall frame-wise displacement (FD) of >0.15 mm (based on the Jenkinson method; Jenkinson et al., 2002) for the Wisconsin dataset and Stanford dataset. In the Leipzig dataset in which four rs-fMRI runs were obtained within participants, we removed runs with more than 0.15 mm of mean FD, and participants based on the mean across runs.

In addition, given that FD can influence observed relationships between functional connectivity and behavior (Siegel et al., 2017), we controlled for FD in analyses focused on relationships between functional connectivity and FFMQ scores (see Methods: “Predictive modeling analysis”).

## Functional connectivity feature extraction

For each individual, we extracted the preprocessed BOLD time series from the mean across all voxels within each node defined based on an intrinsic functional network atlas in MNI space, specifically, the Shen atlas of 268 whole-brain regions (Shen et al., 2013). This atlas has been frequently used in CPM studies (e.g., Kucyi et al., 2021; Rosenberg et al., 2015). We computed Fisher z-transformed Pearson correlation coefficient of time series, giving a matrix of functional connectivity values between all region pairs. We define region pairs as connections or edges.

## Predictive modeling analysis

We chose the Wisconsin sample for training as it is the largest of the datasets in this study (Poldrack et al., 2020). We performed CPM using publicly available code (https://github.com/DynamicBrainMind/CPM_CONN). For each participant in the Wisconsin sample, we generated model-based predictions of FFMQ or FFMQ subscales based on data from all other included participants, i.e., leave-one-participant-out cross-validation (LOOCV)). In each cross-validation fold, we computed the Pearson correlation between each unique edge (pair of nodes) in the functional connectivity matrix (derived from the Shen atlas) and participant FFMQ scores. We then ‘masked’ the brain-behavior correlations: retaining only edges positively or negatively correlated with FFMQ scores at the suprathreshold level of P < 0.01 (two-tailed). This resulted in positive and negative edge masks. For each participant, we computed the dot product between the functional connectivity matrix and each mask. We then calculated a single network strength value, the subtraction of negative edge from positive edge sums. Finally, we fit a linear model, based on all participants within the fold, of the form:

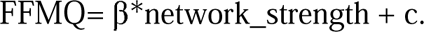

In order to assess whether predicted versus observed scores in LOOCV held-out participants were statistically significant at the group level, we generated a distribution of null values. To do so, we repeated all of the described CPM procedures, except the participant assignments of the FFMQ scores were randomly permuted (1000 iterations) to generate null correlation values. To compute a *p-*value, we then calculated the probability of finding a null correlation at or above the true correlation (predicted versus observed FFMQ).

We repeated CPM procedures controlling for head motion by calculating partial correlations (*partialcorr* in MATLAB) between predicted and observed FFMQ. Using these partial correlations, we controlled for head motion, defined as the mean FD value per participant. We did not conduct permutations for the partial correlation tests because effect sizes were comparable to those obtained in the main analyses. We also controlled for participant asthma status by assessing partial correlations. Finally, we repeated CPM procedures with 10-fold CV, which is a preferred approach for sample sizes > 100 (Poldrack et al., 2020).

## Model selection

We trained models for each of the seven FFMQ scores (Total, Total w/o Observe, and 5 subscales). Models were selected for generalizability testing based on an uncorrected *p* threshold < 0.05 based on the permutation testing (for a similar approach, see Kim et al., 2023).

Our main subsequent analyses focus only on the selected models. For external validation analyses assessing generalizability in the other dataset, as well as analyses of edge network identities, we computed CPM parameters and positive and negative masks based on data from *all* participants in the Wisconsin sample (i.e., a single fold).

## Validation in test samples

We took the selected trained models and applied them on the Leipzig resting-state data (averaged across runs, excluding runs with head motion, as described previously) and Stanford resting-state data (excluding participants with head motion). Each functional connectivity matrix was masked with the positive and negative masks, and then those FC values from the selected edges were either applied independently or summed to form a network strength value. The connectivity predictor or network strength was then used in the linear model to predict the FFMQ scores. We compared FFMQ predicted and observed values using Pearson’s correlation. We also conducted partial correlations controlling for FD values.

## Test-retest stability

There are some indications that CPM predictions may be more reliable and stable than individual edges (Taxali et al., 2021). To test this, we leveraged the Leipzig dataset, which had multiple 15-minute runs. We examined Pearson’s correlations between the CPM network strengths for the first two runs and the last two runs. Additionally, we randomly selected 1000 individual edges, and estimated the probability of the CPM correlation compared to the distribution of random edges.

## Analysis of functional connectivity patterns contributing to Mindfulness CPMs

To gain insight into the neuroanatomical patterns that contributed to the CPMs, we examined brain networks, nodes, and regions. The principal measure for display is ‘degree’, where a high degree means that a node/network/region is involved in many edges. First, we used the WASHU network labels to assign nodes to ten networks (Power et al., 2011). The WASHU networks consist of SMN: somatomotor network, CO: cingular-opercular network, AUD: auditory network, DMN: default-mode network, VIS: visual network, FPN: frontoparietal network, SAL: salience network, SUB: subcortical network, VAN: ventral attention network, and DAN: dorsal attention network. A proportion of the nodes are not assigned to a network. Despite this limitation, the WASHU network labels were chosen because they include networks of interest (FPN, DMN, SAL, SMN, and VIS). We examined the number of connections within each network in matrix plots. Further, we plotted the specific node ‘degrees’ on the brain medial and lateral surfaces using BioImage Suite (https://bioimagesuiteweb.github.io/webapp/connviewer.html). Finally, connectograms were plotted to display connections between brain regions using BioImage Suite.

Comparisons of models were assessed, specifically the overlaps between edges selected by the models. We conducted non-parametric permutation tests (shuffling the edges) to assess whether the degree of overlap was higher than chance.

## Exploratory analysis

We conducted exploratory analyses to identify whether different analysis approaches lead to improvements in generalizability. First, we considered whether generalization to dataset acquired with a different MRI scanner type may be a high bar. Thus we combined the data across scanners before conducting training-test splits. In one analysis of this combined data, we conducted partial correlations using the means of the FFMQ within each dataset to control for dataset differences. Second, we examined whether other predictive modeling methods previously used in neuroimaging applications were more powerful than CPM, including tangent parameterization of connectivity, Brain Basis Sets and Elastic Net Regression. See **Supplement** for Full Methods.

## Results

### Behavioral measures

The mean FFMQ in the Wisconsin sample was 134.8 (*SD* = 17.8), in the Stanford sample 126.8 (*SD* = 17.4), and the in the Leipzig sample 107.2 (*SD* = 11.3) (**Supplementary Figure 1**). Wisconsin FFMQ scores were significantly higher than Stanford scores as assessed by a two-tailed Welch’s t-test (*t*(153.46) = 3.50, *p <* .001, Cohen’s *d* = 0.44), and Leipzig’s (*t*(221.96) = 15.45, *p <* .001, Cohen’s *d =* 1.35). Stanford FFMQ scores were significantly higher than Leipzig FFMQ scores (*t*(139.68) *=* 8.49, *p <* .001, Cohen’s *d =* 1.10). This difference in the trait measures could be related to the student sample of the Stanford dataset, or the German translation of the FFMQ, or cultural differences. FFMQ scores may vary across different samples (e.g., (Goldberg et al., 2016; Isbel et al., 2020). Correlations between the subscales were significant but were all less than *r* = .5, indicating some independence (see **Supplementary Table 1**). Subscales correlated with the total FFMQ score, *rs* > .6, and showed differences across sites similar to total FFMQ score differences (**Supplementary Figure 2** and **Supplementary Figure 3)**.

## Learning the neural features from the training dataset

We trained seven CPMs in the Wisconsin dataset, one for each subscale, the total score, and the total without observing. 18 participants were removed due to above-threshold head motion. Full training set performance is reported in **Supplementary Table 2**. The models predicting *Acting with Awareness* (AA), and *Non-judging* (NJ) showed positive correlations (**Figure 1**) between overall network strength (positive network - negative network) and the respective subscale (AA: *r(*186) = .22, NJ: *r(*186) = .21). The two models had non-parametric *p* values of .017 and .025, respectively, so we selected them for model testing in held out data. When using 10-fold cross-validation instead of leave-one-out cross-validation (LOOCV), the results were similar (AA: *r*(186) = .16, *p* = .046; NJ: *r*(186) = .22, *p =* .021). The single-fold AA model, which we call the AA-CPM, consisted of 328 positive edges and 758 negative edges. Positive edges were present in 95.0% of LOOCV folds, and negative edges were present in 93.0% of LOOCV folds. Partial correlations with framewise-displacement as a covariate showed similar effect sizes (*r*(186) = .22), indicating no influence of head motion on model prediction. Partial correlations with asthma status likewise resulted in similar effect sizes (*r*(186) = .21). The single-fold NJ model, which we call the NJ-CPM, consisted of 664 positive edges and 628 negative edges. Positive edges were present in 94.3% of LOOCV folds, and negative edges were present in 93.6% of LOOCV folds. Partial correlations with framewise-displacement and asthma status as covariates were similar (*r*s of .21 and 0.19, respectively).

**Figure 1:**
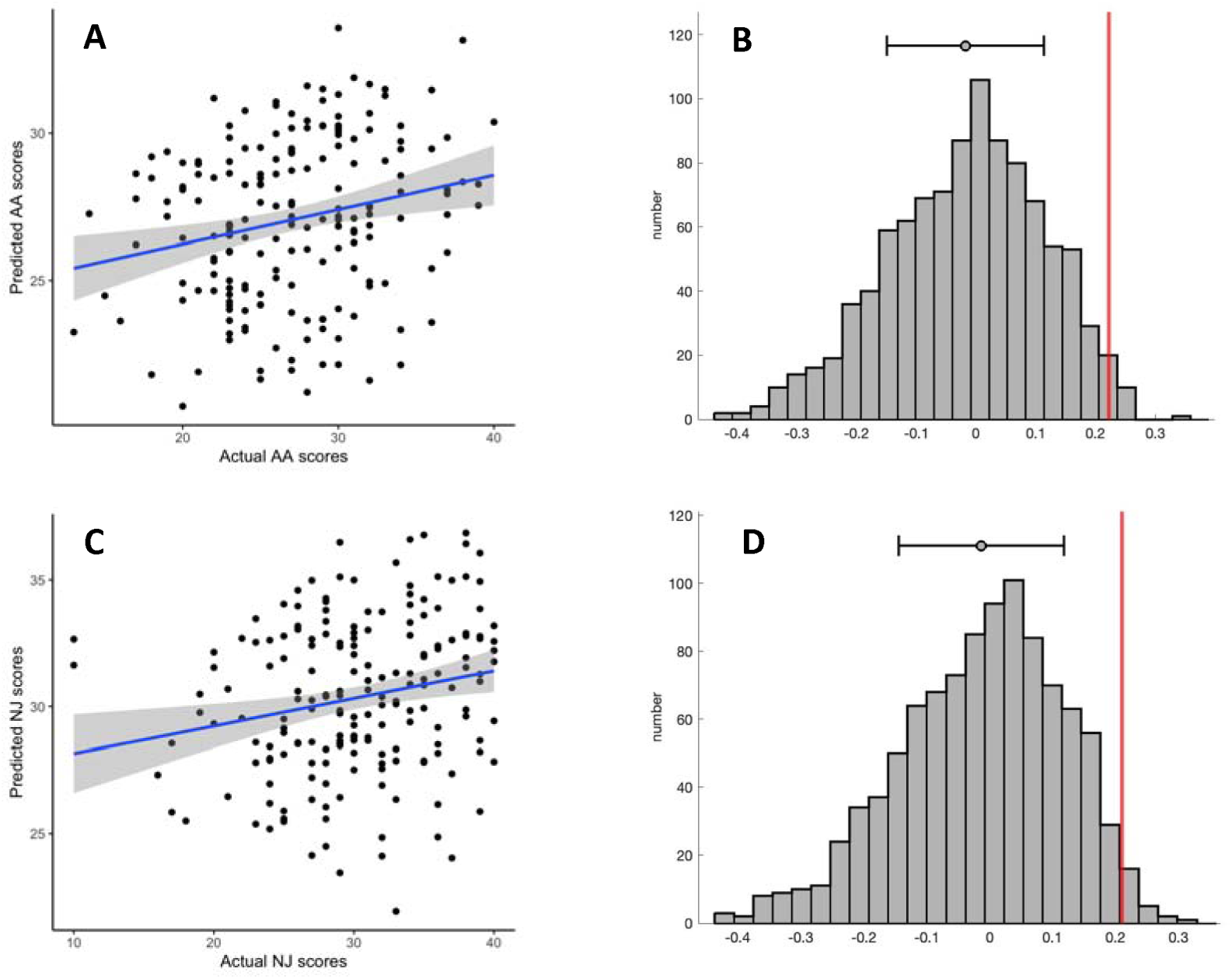
Prediction performance in the training set. A: Predicted vs observed values from leave-one-out cross validation, for *Acting with Awareness* subscale. B: Correlation coefficient (in red) compared to the distribution of null correlation coefficients, for *Acting with Awareness* subscale. C: Predicted vs observed values from leave-one-out cross validation, for *Non-judging* subscale. D: Correlation coefficient (in red) compared to the distribution of null correlation coefficients, for *Non-judging* subscale. In plots A and C, grey shading reflects 95% confidence intervals. In plots B and C, the mean and standard deviations are shown above the distributions.

## Features in the AA-CPM

We next analyzed the masked edges from the AA model, as derived from a single fold (**Figure 2**). In the positive network, FPN between-network connections were most featured, primarily between FPN and sensory networks as well as FPN-DMN. There were some connections incorporating DMN, SAL and SMN. High-degree brain areas included the cerebellum, parietal areas, and dorsomedial prefrontal cortex. Circle plots from the FPN, in particular, show mostly cross-hemispheric connections between prefrontal, motor areas, parietal areas, and limbic areas, with more diverse brain areas in the right hemisphere (**Supplementary Figure 4)**. The negative network (edges that negatively correlated with AA scores) contained connections within the SMN, and between the SMN and the VIS network, with some involvement of auditory and DMN networks (**Supplementary Figure 5)**. High degree nodes included somatomotor cortices, primary occipital cortices, and ventrolateral prefrontal cortex. Circle plots demonstrated dense connections across hemispheres between motor, parietal, temporal, visual, and insula areas (**Figure 3)**.

**Figure 2:**
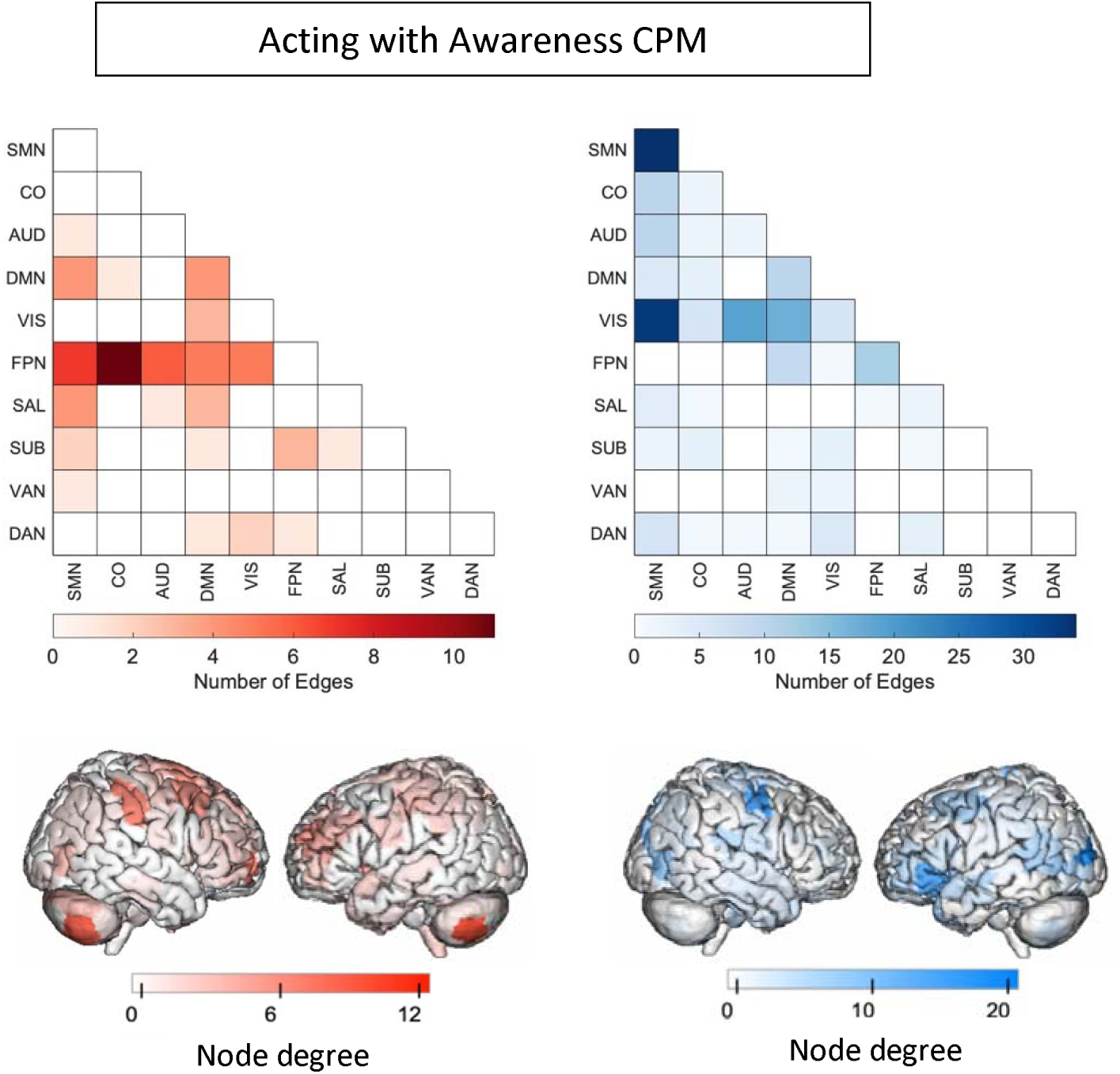
Edges included in single-fold model of *Acting with Awareness* subscale of FFMQ using Shen atlas. In red are edges that positively predict *Acting with Awareness*. In blue are edges that negatively predict *Acting with Awareness*. Node degree: number of connections including that node (brain area). SMN: somatomotor network, CO: cingular-opercular network, AUD: auditory network, DMN: default-mode network, VIS: visual network, FPN: frontoparietal network, SAL: salience network, SUB: subcortical network, VAN: ventral attention network, DAN: dorsal attention network.

**Figure 3.**
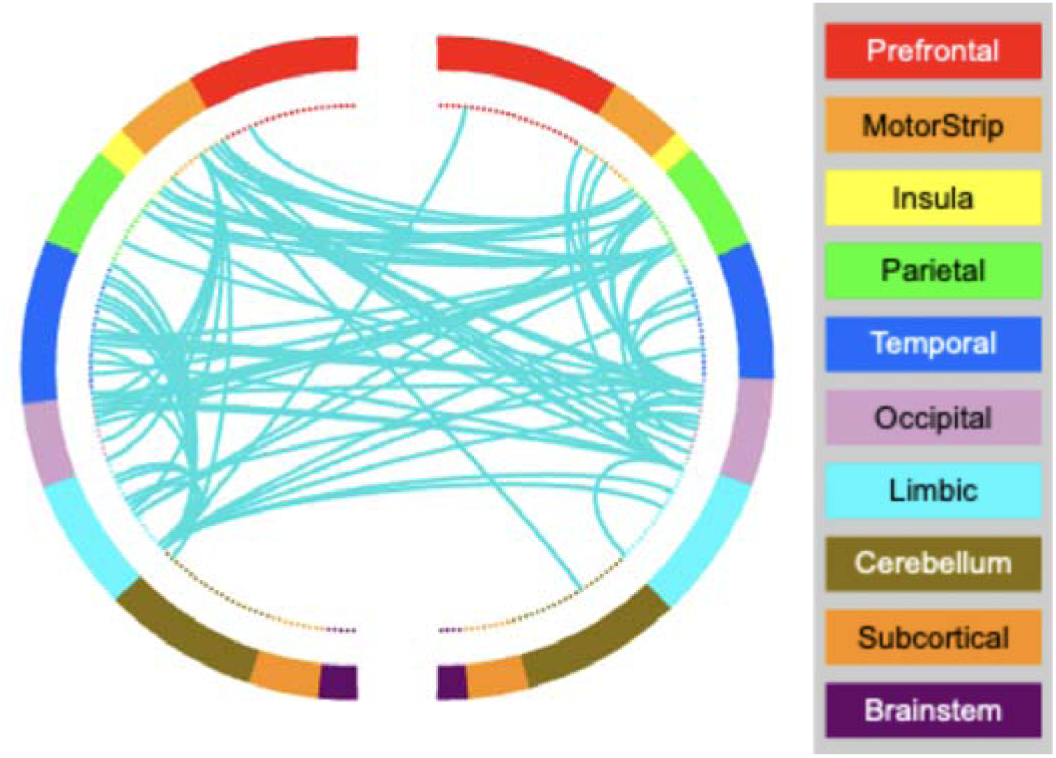
AA-CPM Connectogram. Negative network, degree threshold = 15.

## Features in the NJ-CPM

We analyzed the masked edges from the NJ model derived from a single fold (**Figure 4**). In the positive network, DMN connections to the rest of the brain were most featured, with some additional connections involving FPN and SUB. DMN-SMN and DMN-CO edges were most prevalent. High degree brain areas included the ventromedial prefrontal cortex, posterior cingulate cortex, and medial somatomotor areas. Circle plots of DMN connections show dense connections between left limbic areas and left motor areas, as well as the insula and parietal areas (**Supplementary Figure 6)**. The negative network (edges that negatively correlated with NJ scores) was widely distributed, with many edges in the SMN network, and between VIS and DMN (**Supplementary Figure 7**). High degree brain areas included bilateral occipital areas, posterior temporal lobe including temporoparietal junction, and bilateral somatomotor areas. Circle plots demonstrated similar connections to the AA-CPM negative network, with the exception of right subcortical involvement (**Figure 5**).

**Figure 4:**
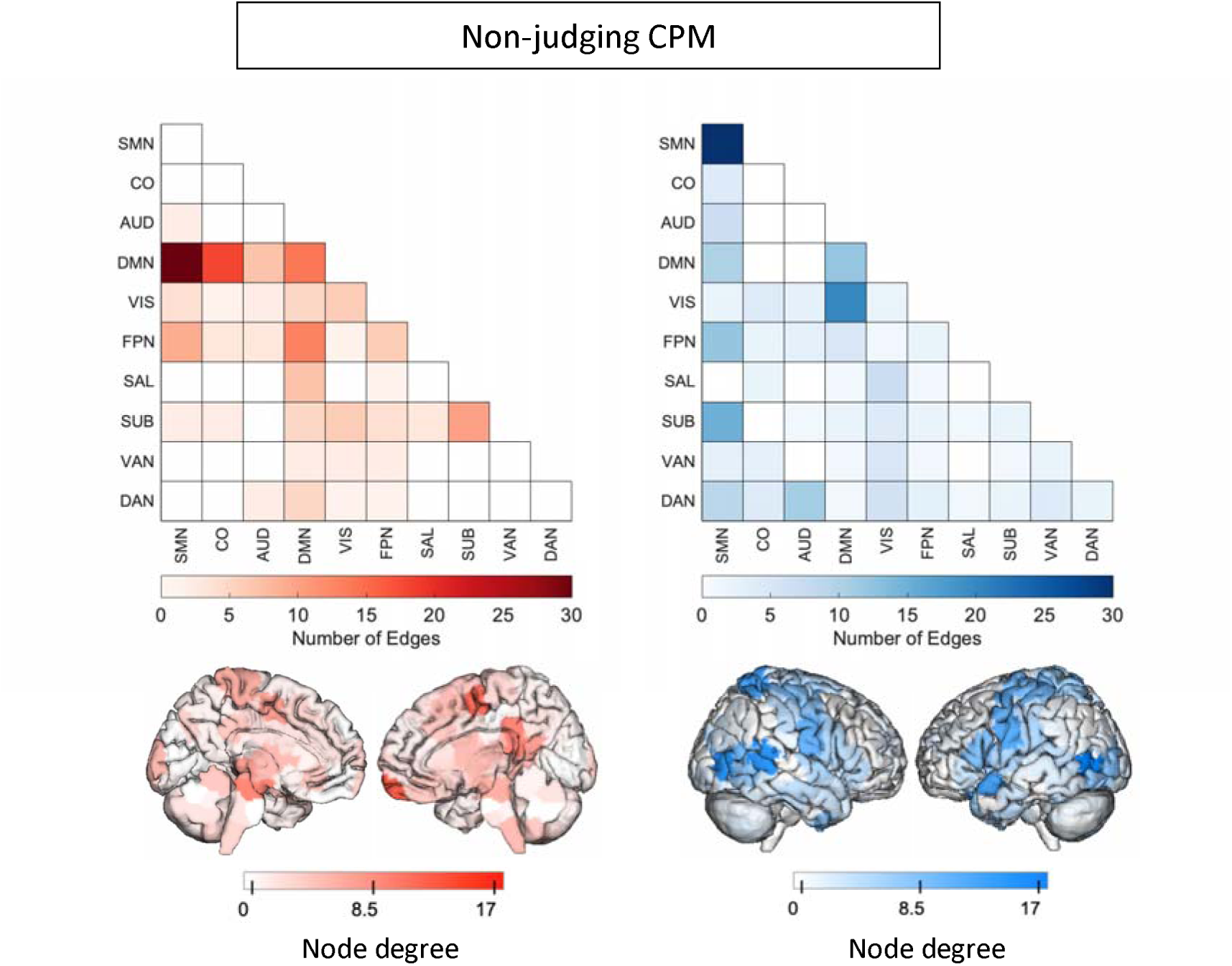
Edges included in single-fold model of *Non-judging* subscale of FFMQ using Shen atlas. In red are edges that positively predict *Non-judging*. In blue are edges that negatively predict *Non-judging*. Node degree: number of connections including that node (brain area). Medial views are shown for positive network as high-degree nodes are medial. SMN: somatomotor network, CO: cingular-opercular network, AUD: auditory network, DMN: default-mode network, VIS: visual network, FPN: frontoparietal network, SAL: salience network, SUB: subcortical network, VAN: ventral attention network, DAN: dorsal attention network.

**Figure 5.**
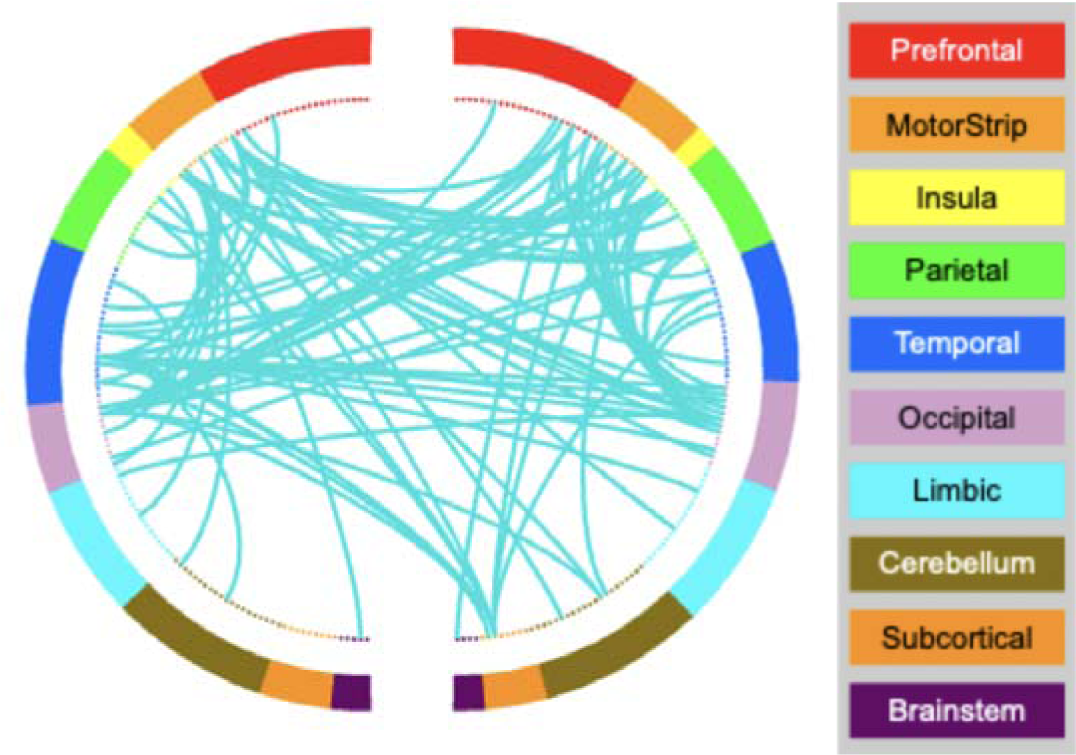
NJ-CPM Connectogram. Negative network, degree threshold = 15.

## Testing performance of AA-CPM

When we applied the AA-CPM to the Leipzig dataset, we found a significant positive association between predicted and observed scores (*r*(75) = .34, *p* = .0025, two participants were removed due to head motion) (**Figure 6A**). The association remained significant when partialling out head motion (*pr*(75) = .27, *p* = .020). The positive association was significant when examining only the negative network (*r*(75) = .33, *p* = .0031), but not the positive network (*r*(75) = .032, *p* = .78). We found no significant relationship between AA-CPM predictions (positive, negative, and network strength) and observed AA scores in the Stanford dataset (*ps* > 0.5).

**Figure 6.**
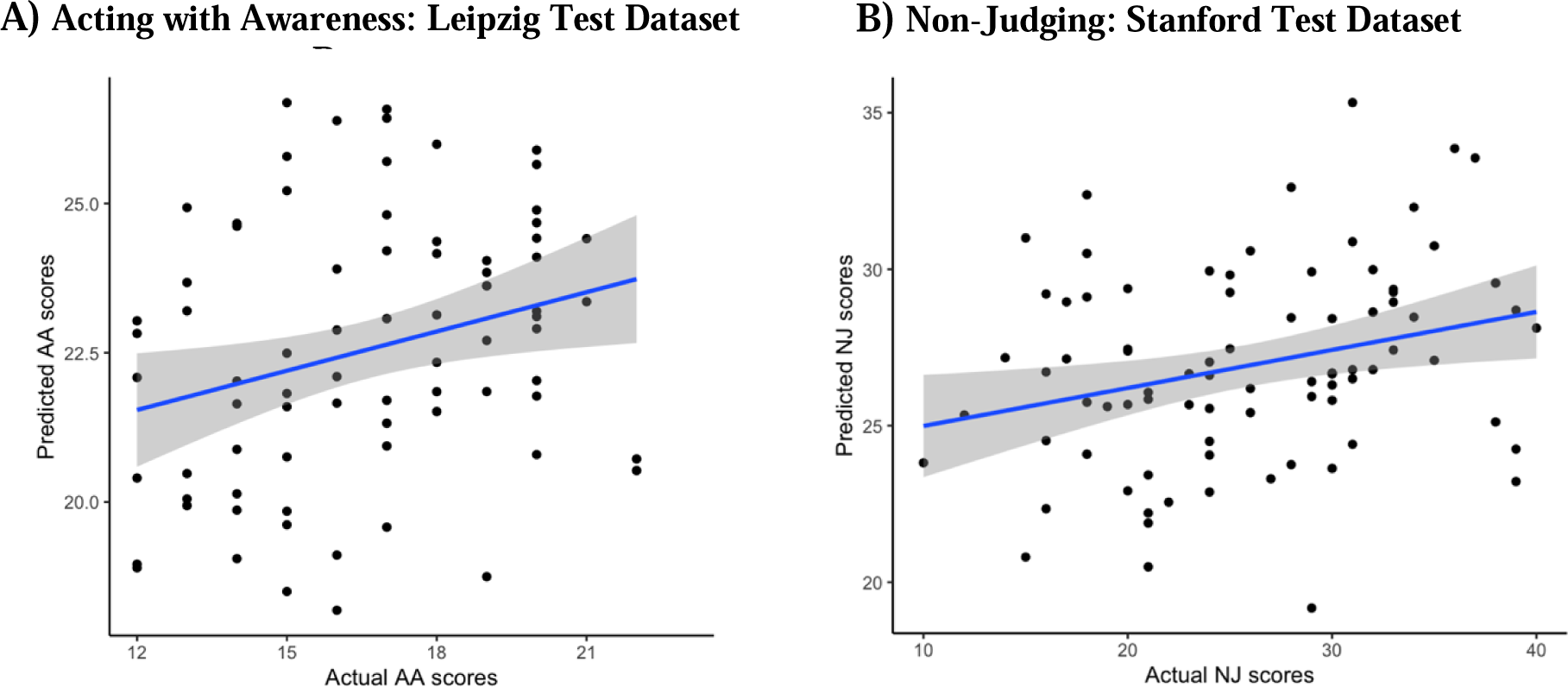
Test predictions vs. Actual scores. A) Leipzig held-out data prediction performance, for AA-CPM predicting AA scores. Blue line is linear best fit, with gray 95% confidence intervals. B) Stanford held-out data prediction performance, for NJ-CPM predicting NJ scores. Blue line is linear best fit, with gray 95% confidence intervals.

## Testing performance of NJ-CPM

When we applied the NJ-CPM to the Leipzig dataset, we found no association between predicted and observed scores (*ps >* 0.2, two participants were removed due to head motion). We found a positive relationship between NJ-CPM predictions and observed NJ scores in the Stanford dataset (*r*(80) = .28, *p* = .012) (**Figure 6B).** The positive association was still present when just examining the negative network (*r*(80) = .27, *p* = .016), but not the positive network (*r*(80) = .16, *p* = .16). Results were identical when partialling out head motion.

## Other model validation steps

### Relationships between AA-CPM and NJ-CPM

NJ and AA subscale scores were positively correlated in the Wisconsin training dataset (*r(*204) = .36, *p* < .001). In addition, there was a strong positive correlation between the AA-CPM and the NJ-CPM strengths across all datasets, (*r*(345) = .66, *p* < .001). 20 positive edges were shared between the models (2.02%, non-parametric *p* < 0.001), and 58 negative edges were shared between the models (4.18%, non-parametric *p* < 0.001). To evaluate the specificity of the AA-CPM and NJ-CPM, we examined whether they could cross-predict in the held-out datasets. The NJ-CPM could predict AA in the Leipzig dataset (*r*(75) = .29, *p* = .0089). The AA-CPM trended towards predicting NJ in the Stanford dataset (*r(*80) = .20, *p* = .065). Results were similar when just examining the negative networks. As a control, we examined whether a non-significant model from the Wisconsin dataset (the Observing-CPM), could predict in the hold out data. The Observing-CPM did not predict AA in the Leipzig dataset (*r*(75) = .098, *p* = .39), nor NJ in the Stanford dataset (*r*(80) *=* .14, *p* = .22).

### Relationships with mind-wandering CPM

Trait mindfulness has been found to be negatively correlated with mind-wandering (MW) on tasks (Belardi et al., 2021; Mrazek et al., 2012). We examined the correlation between network strengths for the AA-CPM and NJ-CPM and a previously published CPM for mind-wandering, the MW-CPM (Kucyi et al., 2021), in data from all sites. There was a negative correlation between AA-CPM and MW-CPM, (*r*(345) = -.22, *p* < 0.001) (**Supplementary Figure 12**), as well as between NJ-CPM and MW-CPM (*r*(345) = -.25, *p* < 0.001) (**Supplementary Figure 13**). Overlaps between edges for AA-CPM vs MW-CPM and NJ-CPM vs MW-CPM were not significantly higher than chance (non-parametric *p >* 0.1). The In the held out datasets, MW-CPM negatively predicted NJ in the Stanford dataset (*r*(80) = - 0.31, *p* = 0.0042), and no other predictions were statistically significant (*ps* > 0.5).

### Test-retest stability

We also examined the test-retest stability of the models in the Leipzig dataset, by comparing the network strengths for the first two runs to the last two runs using Pearson’s correlations. The AA-CPM and NJ-CPM showed correlations (*r*(74) = .35, *r*(74) = .41) that were not more stable than random edges (non-parametric *ps* > 0.5).

## Exploratory analyses

In the combined, shuffled data, we found a significant model predicting NJ, which generalized to the held-out dataset. It should be noted that the positive network for the supplementary NJ model was similar to the positive network found in the main NJ-CPM. Results from the tangent parameterization of connectivity, Brain Basis Set regression, and Elastic Net Regression failed to generalize from Wisconsin to other datasets.

## Discussion

We used connectome predictive modeling (CPM) to investigate the relationships between trait mindfulness as measured by FFMQ, and functional networks across the whole brain (including the DMN, FPN, and SN) and assessed whether the relationships generalized to independent samples. With 367 participants over three sites, this is the largest neuroimaging study of trait mindfulness to-date (Treves et al., in press). While we did not find a generalizable model of total FFMQ scores, we did find models of the *Acting with Awareness* and *Non-judging* subscales that generalized to one of two independent datasets. The models showed highly similar negative networks (i.e., increasing connectivity negatively predicts mindfulness) involving DMN, VIS and SMN connectivity, and these negative networks were responsible for generalization performance. Our findings highlight the importance of networks not investigated previously (e.g. VIS, SMN) (Treves et al., in press), inform new frameworks for defining trait mindfulness (Altgassen et al., 2023), and underscore the difficulty of using neuroimaging measures for predicting individual differences (Marek et al., 2022).

## Networks implicated in trait mindfulness

Previous studies on trait mindfulness have focused on seed-based analysis of the triple networks: the DMN, FPN, and SN. A large cognitive neuroscience literature has implicated these networks in the regulation of external and internal attention (Buckner & DiNicola, 2019; Menon, 2011). In keeping with this, meta-analytic reviews have found that mindfulness interventions lead to increases in DMN-SN connectivity (Rahrig et al., 2022) and mindfulness practice (focused attention) leads to decreased activations in DMN regions like the posterior cingulate cortex compared to control conditions (Ganesan et al., 2022). Despite this, there is no consensus with regards to the networks’ relationships to trait mindfulness. There are some indications of triple network involvement in predicting trait mindfulness in the current study. The predictive models developed here consist of positive networks (connectivity that predicts higher trait mindfulness) and negative networks (connectivity that predicts lower trait mindfulness). It is important to note that the models consist of hundreds of edges across the entire brain and here we summarize notable networks and regions. The positive network for the *Acting with Awareness* model included connections between FPN and other brain networks including sensory networks and DMN, with high-degree nodes (regions involved in many connections) in the cerebellum, dorsomedial prefrontal cortex, and parietal cortex. FPN connectivity could be related to top-down regulation of attention (Marek & Dosenbach, 2018), and individuals who score high on *Acting with Awareness* may regulate their attention using the FPN. However, this connectivity did not generalize to predict scores in the test datasets. Instead, connections involving the DMN, SMN, and VIS networks negatively predicted mindfulness scores (both *Acting with Awareness* and *Non-judging)* in the training and test datasets (and under different training-test splits of the data). Decreases in DMN connectivity with the rest of the brain could reflect differences in habitual self-referential processing, e.g. rumination (Butterfield et al., 2023; Frewen et al., 2020; Raichle et al., 2001; Zhou et al., 2020). The somatomotor (SMN) network has been implicated in mind-wandering (Kucyi et al., 2024; Mckeown et al., 2020; Vatansever et al., 2019). Speculatively, altered SMN connectivity may reflect different habitual processing of afferent thermo-ceptive, proprioceptive or even pain signals, or it may reflect simulated motor action (Sormaz et al., 2018). Associations with visual network connectivity may reflect differences in sensory awareness, and visual network connectivity has been observed to change after mindfulness training (Kilpatrick et al., 2011).

Evidence that the models capture meaningful neural function is that they were significantly negatively correlated with a well-established mind-wandering CPM (Kucyi et al., 2021). This is despite not involving the same brain connections. Mind-wandering and mindfulness may be thought of as intuitive opposites, e.g. participants rating high on mindfulness questionnaires mind-wander less (Mrazek et al., 2012). Notably, the mind-wandering model was trained on state mind-wandering ratings during an attention task, not trait questionnaires. Perhaps the mindfulness models developed here could predict mindful states as well as traits.

## Distinctions between mindfulness scales

The models that met our selection threshold in the training dataset were trained on *Acting with Awareness* and *Non-judging.* These two facets of mindfulness make up attitudinal and attentional components of mindfulness (Rau & Williams, 2016) and are common in survey instruments measuring mindfulness (Altgassen et al., 2023). The *Acting with Awareness* subscale of the FFMQ is typically thought to reflect an attentional component of mindfulness (Baer et al., 2006). It involves questions from the Mindful Attention and Awareness Scale (MAAS) (Brown & Ryan, 2003), including “It seems I am running on automatic without much awareness of what I’m doing” (reverse-coded). A network analysis found that *Acting with Awareness* clusters with the MAAS, mind-wandering questionnaires, and cognitive failures questionnaires (Beloborodova & Brown, 2023). Finally, scores on *Acting with Awareness,* and the MAAS, respectively, correlate with objective behavioral measures of attention (Ching & Lim, 2023; Mrazek et al., 2012). Our study provides more external validation of self-report mindful attention by identifying resting-state connections across the whole brain that predicted *Acting with Awareness* scores.

The *Non-judging* subscale of the FFMQ is typically thought to reflect affective components of mindfulness, specifically one’s tendency to become aware of thoughts and feelings without judgement (Baer et al., 2006). An example item is “I criticize myself for having irrational or inappropriate emotions” (reverse coded). Correlations between positive mental health outcomes and *Non-judging* are often found (Blanke et al., 2018; Cortazar & Calvete, 2019; Treves et al., 2023). One theorized link is through decreased rumination (Greco et al., 2011), or perseverating on negative self-referential thoughts, memories, and one’s own negative mood (Mennin & Fresco, 2013; Nolen-Hoeksema, 1991).

One area of uncertainty highlighted by the current study is whether *Acting with Awareness* and *Non-judging* are distinct or whether they correspond to a single ontological concept of mindfulness. Standard definitions of mindfulness often unite attentional and attitudinal features of mindfulness, e.g. mindfulness is a present-focused attention, with an orientation of acceptance and non-judgement (Bishop et al., 2004). Empirically however, this unity may not be supported. More recent research on self-report mindfulness has indicated that single-factor definitions of mindfulness may be neither accurate nor predictive of real-world outcomes (Altgassen et al., 2023; Bednar et al., 2020; Beloborodova & Brown, 2023; Tran et al., 2020). Our study contributes to this debate but does not resolve it. In the large training sample, there were discriminable brain connections (e.g., FPN vs DMN) that positively predicted *Acting with Awareness* and *Non-judging.* However, the brain connections that predicted these subscales in the relatively smaller held-out datasets were largely overlapping. It may be that large datasets are necessary to identify their distinctions neurally. It is also unclear why other subscales and the total FFMQ scores were not predictable. One possibility is that the shared variance between *Acting with Awareness* and *Non-judging* may be more predictable neurally than the FFMQ total scores which combine across 39 distinct items. Subscales like *Observing* are sometimes left out from measurement because they are understood differently by different populations (Baer et al., 2022; Gu et al., 2016; Pang & Ruch, 2019). An important consideration is that introspective ability confounds self-reported mindfulness measurement (Grossman, 2011), and more recent research has attempted to control for the reliability of responders before conducting correlations with functional neuroimaging measures (Y. Kim et al., 2023). To summarize, our results speak to the possibility of different operationalizations of mindfulness measurement – operationalizations for insight into self-reported experiences may differ from operationalizations that align with objective measurement.

## Limitations and the difficulty of individual differences in neuroimaging research

We did not find complete model generalization. The negative network models (involving DMN, SMN, and VIS) predicted *Acting with Awareness* in the Leipzig dataset, and *Non-judging* in the Stanford dataset. One possibility is that the cross-site differences were a high barrier to generalization. The data acquisition parameters varied from site to site. Scanner parameters have an impact on activations and connectivity estimates (Friedman et al., 2008; Glover et al., 2012; Greve et al., 2013). In addition, the trait mindfulness scores varied significantly from site to site, and the connections that predict mindfulness in one range of scores may not generalize to another range. It is unclear whether overall score differences between sites reflect real individual differences (assumption 1) or noise (assumption 2) (in which case they should be z-scored within site). In exploratory analyses, we tested the robustness of our models to this assumption, and the negative network models still showed significant prediction performance in training and test splits. To our knowledge, variability in baseline trait mindfulness scores has not been explored, and the ramifications for neuroimaging studies are unclear. Indeed, variability in mindfulness scores may also have been a key factor for the training performance of the models.

A second limitation concerns the reliability/stability of the CPM measures. A previous study found that when examining split-halves of a 30-minute resting-state scan, CPM networks were more reliable than individual edges (although edges showed a wide range of reliabilities with many exceeding that of CPM) (Taxali et al., 2021). We did not find this to be the case in our study. Reliability puts an upper limit on correlations between outcomes (Dubois & Adolphs, 2016; Nunnally, 1970), and could have limited our ability to find meaningful relationships herein. In a recent study on dynamic functional connectivity, we found that only the most reliable brain measures showed significant relationships with trait mindfulness (Treves et al., 2024).

The predictive modeling results present some concerns as well. In addition to CPM, we conducted elastic net regression, Brain Basis Set regression, and a different connectivity parameterization (tangent-space covariance). Although in the Wisconsin model-training dataset the *Acting with Awareness* and *Non-judging* models showed significant correlations, these alternative predictive models did not generalize to the test datasets (nor did models for the other subscales). Our finding that only the CPM approach generalized to independent data highlights the sensitivity of the method while also suggesting some fragility of the brain-behavior relationships. An important caveat of the methods used in our study is that they do not involve any priors over the features used for prediction (a different approach is the network-based statistics prediction toolbox [NBS-Predict], which is biased towards finding connected sets of features; Serin et al., 2021).

A final limitation reflects the real-world implications of these findings. Even though we combined connections across the whole brain for prediction, the amount of variance explained was low (∼4%). This means that using the models for clinical prediction may not be feasible. This has been proposed to be a limitation of neuroimaging, not the models nor specific measures (Marek et al., 2022). Indeed, in the context of classifying individuals with depression, Winter et al. (2024) used multiple modalities of neuroimaging and tested millions of predictive models (with varying hyperparameters), showing a maximum accuracy of 62%. One source of this difficulty may be between-individual variation in neural substrates. It may be the case that fMRI measures of functional connectivity are more powerful for predicting within-individual variation, e.g. fluctuations related to cognition, sleep or arousal (Flournoy et al., 2024; Kucyi et al., 2024) or states of mindfulness vs inattentiveness (Weng et al., 2020).

## Summary

We conducted the largest neuroimaging study of trait mindfulness to-date, with three independent fMRI datasets constituting 367 participants. We have demonstrated that subscales of trait mindfulness are to some degree represented within common whole-brain patterns at rest. Future work could examine the discriminability of the brain representations and their malleability to mindfulness training.

## Ethics statement

Procedures for the Wisconsin study were approved by the Healthy Sciences Institutional Review Board of the University of Wisconsin-Madison. Procedures for the Leipzig study were approved by the ethics committee at the medical faculty of the University of Leipzig (097/15-ff). The Stanford study was approved by the Stanford University Institutional Review Board.

## Data availability statement

Preprocessed data and code will be uploaded to the following link upon publication: https://osf.io/tz86b/.

## Acknowledgements

We thank Patrick Bissett, Yoonji Lee, Russell Poldrack for help with the Stanford Dataset.

---

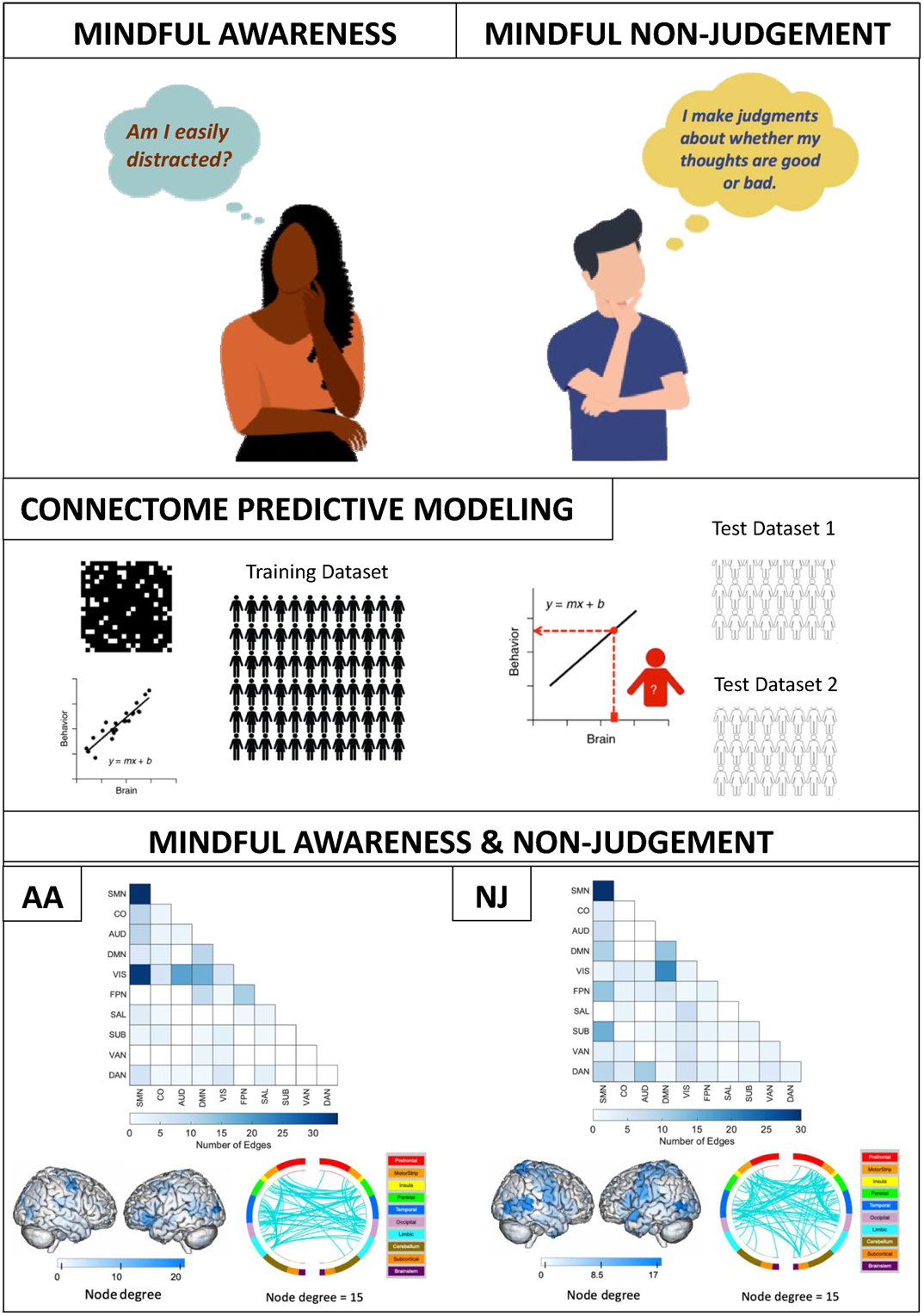

