## Supplement for "Connectome predictive modeling of trait mindfulness"

**Deviations from Preregistration:**

We first published a preregistration upon receiving the Leipzig and Stanford datasets. However, we then were allowed to access the larger NCCAM training dataset. In addition, we were made aware that all datasets contained subscales of trait mindfulness (FFMQ facets). Finally, we decided to test the sensitivity of connectome predictive modelling (CPM) by conducting other predictive models like elastic net regressions. We updated the preregistration on 12/2/2023 to reflect these changes. No models were trained until after updating the preregistration.

Deviations from preregistration:

- We included subjects with clinical diagnoses and asthma (sensitivity to these factors was addressed).
- We did not explore *p*-values for CPM thresholding beyond *p* = 0.01.
- We did not explore atlases besides the 268-node Shen atlas.
- We used a stricter threshold for framewise displacement (mean FD < 0.15 mm) in all samples, but the same scrubbing threshold.
- We examined the validity of our models in the following ways not preregistered:
  - Test-retest stability in the Leipzig sample (with multiple resting-state runs)
  - Cross-prediction of *Acting with Awareness* model, and *Non-judging* model, to assess their independence.

**Exploratory Analysis – Methods:**

***Training splits***

Generalization to another scanner may be a high bar. For this reason, it may be best to combine the data across scanners before conducting training-test splits. This could lead to finding more generalizable features. We assessed this possibility by combining all the data and then conducting an 80-20 train-test split. A limitation of this approach is that training performance may be artificially inflated if the scanner differences are correlated with outcome measures (e.g., the FFMQ). We conducted partial correlations using the means of the FFMQ within each dataset to remove this possibility.

***Predictive modeling***

Other predictive modeling methods may be more reliable across scans than CPM (Taxali et al., 2021) or in some cases provide more prediction accuracy (Dadi et al., 2019). To ensure that results were not limited by the CPM approach, we explored other methods. We examined Brain Basis Sets, which reduces the data (functional connectivity features) using principal component analysis before conducting linear regression of the outcome scores (Sripada et al., 2019). We optimized the dimensionality reduction (n = 25, 50, 75, 100, 125, 150, 175) in the Wisconsin training set. We additionally examined elastic net regression, which penalizes coefficients of the linear regression using two parameters *alpha,* which controls the type of penalty (lasso-ridge), and *lambda,* which controls the extent of the penalty. We optimized these parameters in the Wisconsin dataset. Both approaches were conducted using LOOCV as well as 10-fold CV.

A final innovation was suggested by Dadi et al. (2019), consisting of measuring connectivity differently (Varoquaux et al., 2010). Dadi et al. found optimal phenotype classification performance using tangent-space parameterization of the covariance matrices. We assessed this ­method in conjunction with CPM. Again, as we trained models for each of the seven measures (FFMQ Total, FFMQ Total w/o observe, FFMQ subscales), we controlled for multiple comparisons using FDR-correction (Benjamini & Hochberg, 1995).

***Shuffled datasets***

We shuffled data across the three sites and then split the data into a training set and hold out test set. In the training set (*n* = 271), we achieved prediction performance (when partialling out overall mean score differences between sites) only for the NJ model’s positive network (*r*(269) = .18, *p* = .017). This was not significant when controlling for multiple comparisons (*p_FDR_* = 0.12). This model did generalize to the hold out sample (*r*(71) = .28, *p* = .019). It should be noted that the positive network for the NJ model was similar to the positive network found in the main NJ-CPM (**Supplementary Figure 8**), including DMN-SMN and DMN-CO connections, although many more edges were selected.

***Other predictive models***

Results from the tangent parameterization of connectivity, Brain Basis Set regression, and elastic net regression failed to generalize from Wisconsin to other datasets. For a full reporting of training set performance, see **Supplementary Table 2.**

**Results:**

***Relationships with mind-wandering CPM***

Overlaps between edges for AA-CPM vs MW-CPM and NJ-CPM vs MW-CPM were not significantly higher than chance (non-parametric *p >* 0.1). Further, we examined whether the models had similar features even if the selected edges were distinct, and we found that the networks of FPN, DMN, SMN, and VIS were highlighted by our models and the MW-CPM. However, less negative FPN-DMN correlations were associated with more mind-wandering in their model, and we found the opposite association with ‘acting with awareness’. On the other hand, SMN-VIS associations were both associated with higher mind-wandering and lower scores on ‘acting with awareness’. More positive SMN-DMN correlations were associated with lower mind-wandering and associated with more non-judging in our model. Finally, we observed a strong relationship of SMN-SMN correlations with lower non-judging and lower acting with awareness, but these connections were not implicated in MW-CPM.

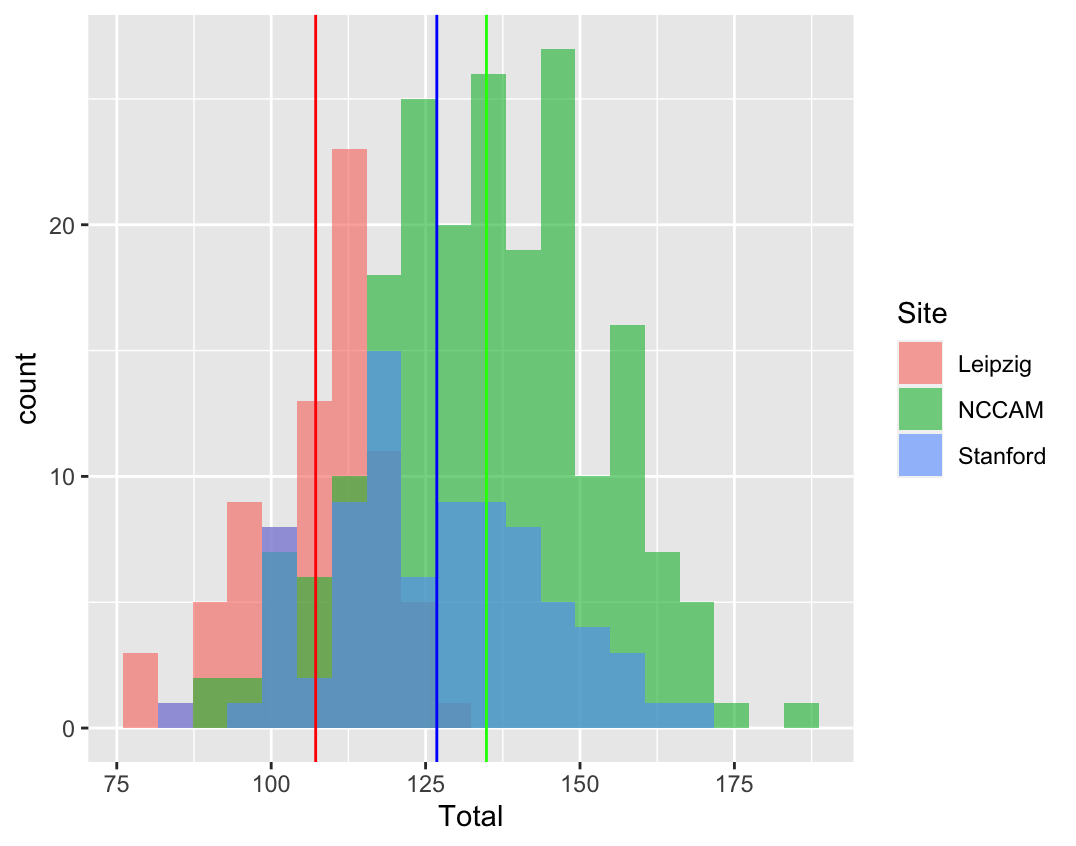

**Supplementary Figure 1.**

Distributions of FFMQ total scores in each dataset. Vertical lines indicate means for each site.

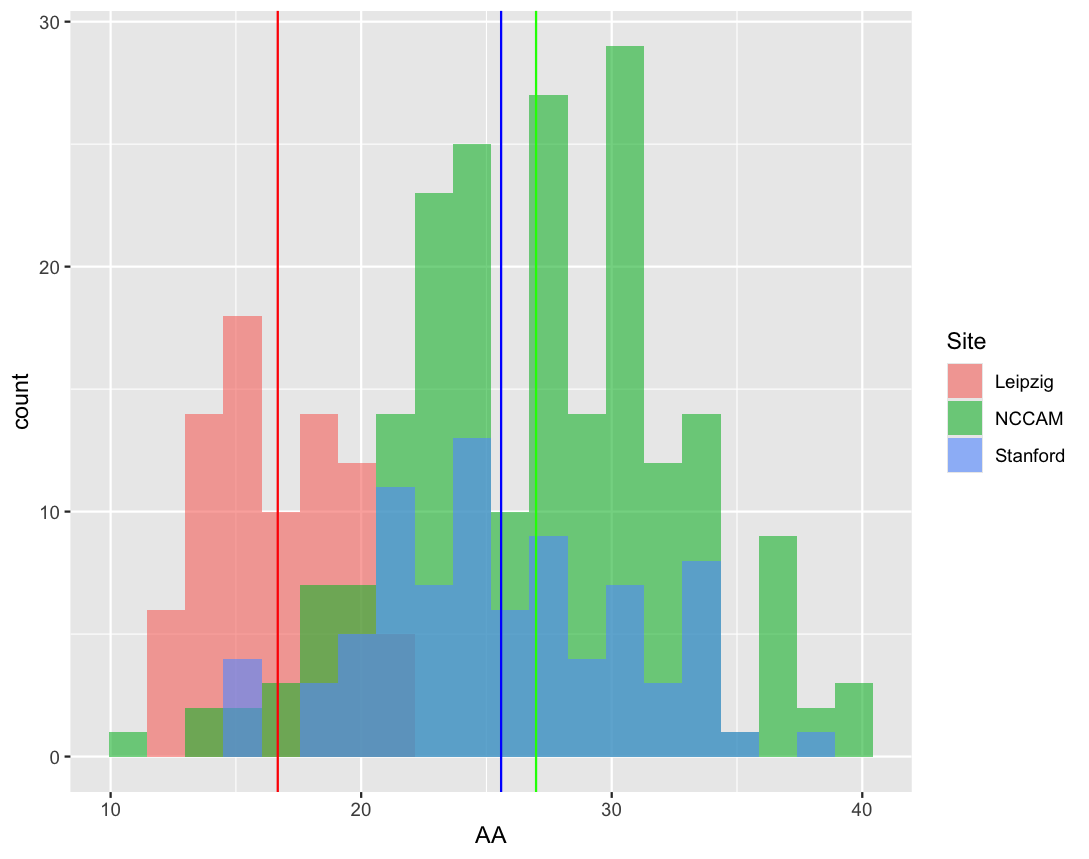

**Supplementary Figure 2.**

Distributions of performance on the Acting with Awareness (AA) subscale. Subscale correlates with total FFMQ score, *rs* > 0.6, and shows differences across sites similar to total FFMQ score differences. Vertical lines indicate means for each site.

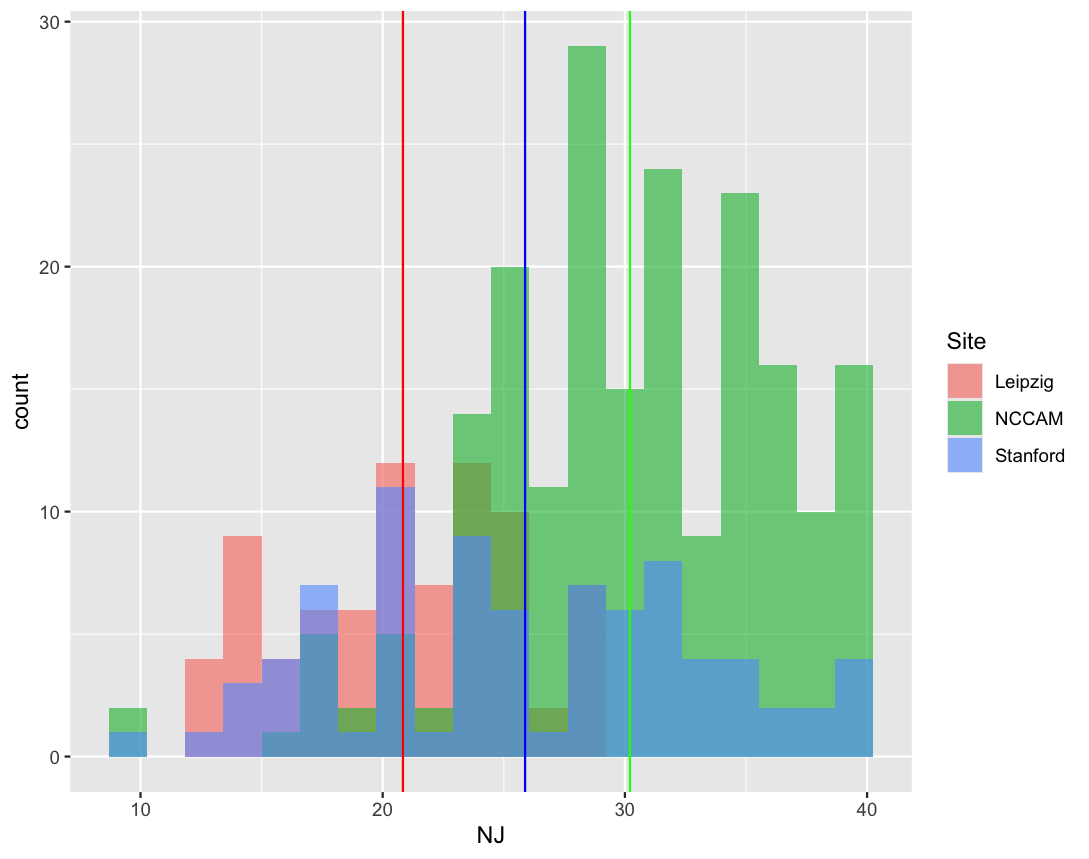

**Supplementary Figure 3.**

Distributions of performance on the Non-judging (NJ) subscale. Subscale correlates with total FFMQ score, *rs* > 0.6, and shows differences across sites similar to total FFMQ score differences. Vertical lines indicate means for each site.

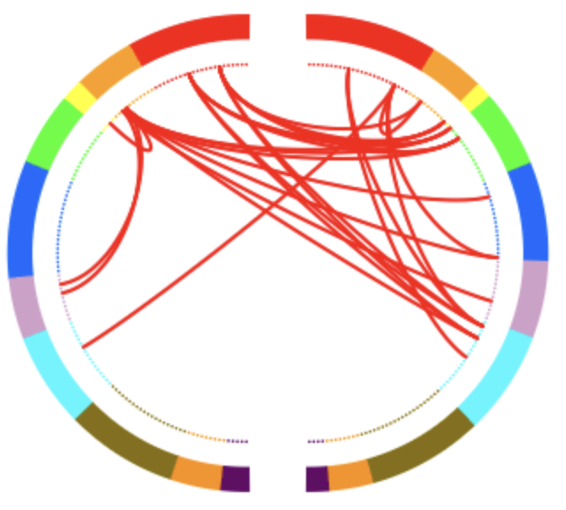

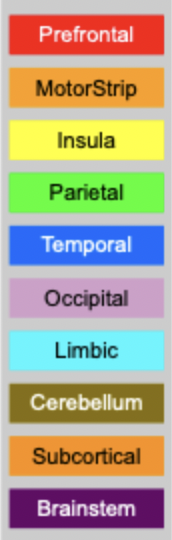

**Supplementary Figure 4.**

AA-CPM Connectogram. Positive network, edges from FPN nodes, degree threshold = 5.

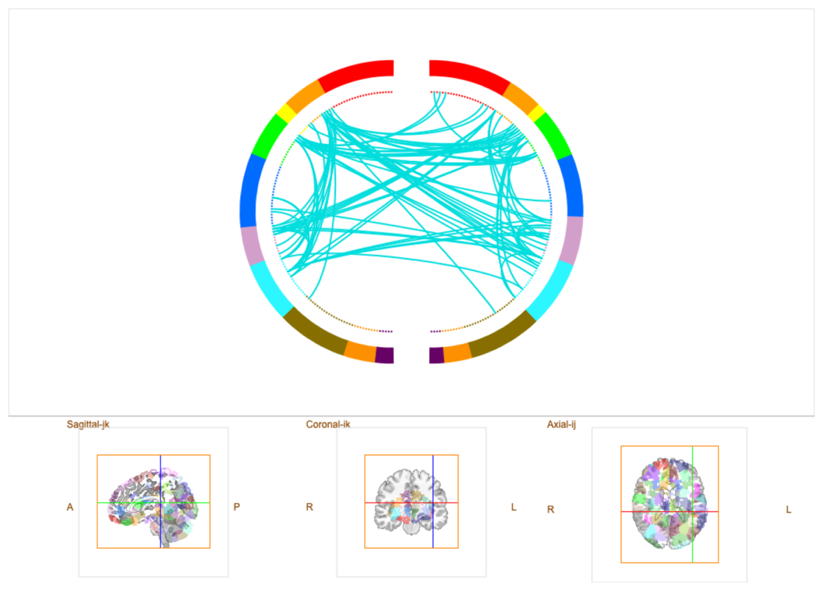

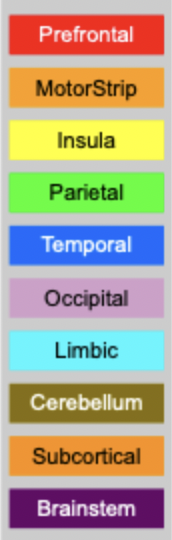

**Supplementary Figure 5.**

AA-CPM Connectogram. Negative network, edges from SMN nodes, degree threshold = 5.

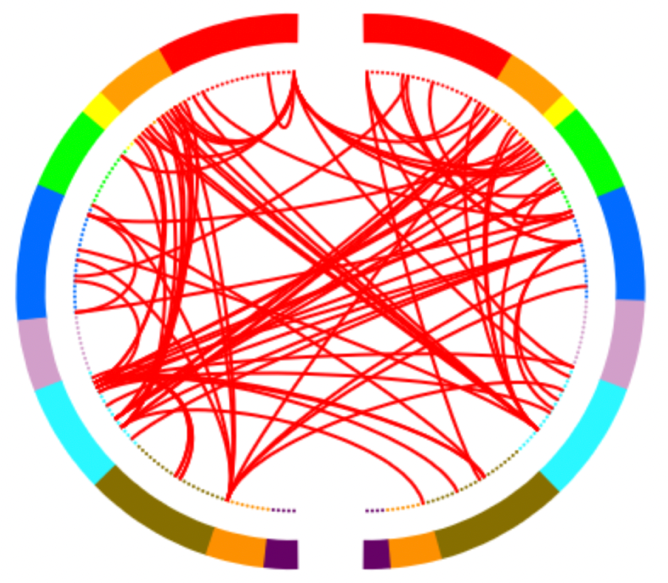

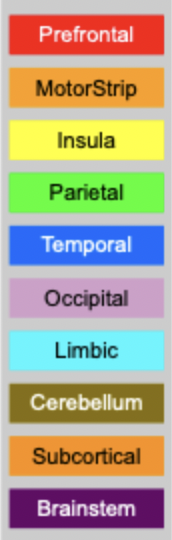

**Supplementary Figure 6.**

NJ-CPM Connectogram. Positive network, edges from DMN nodes, degree threshold = 5.

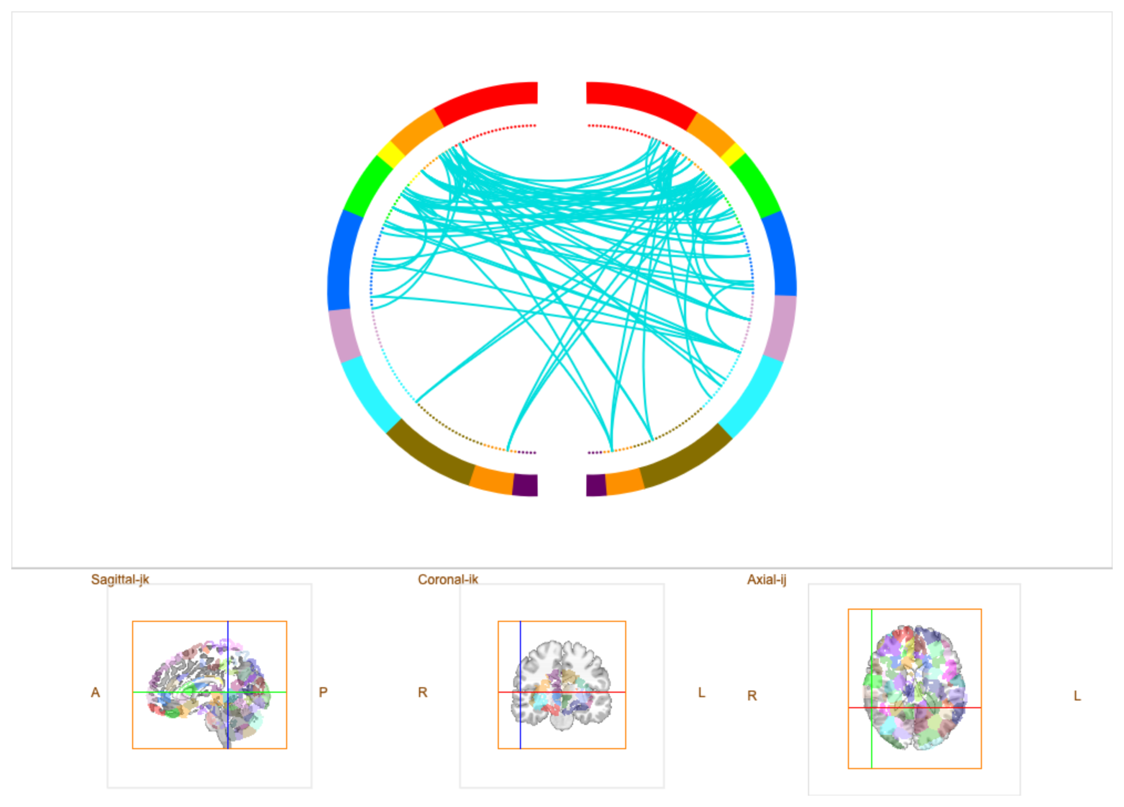

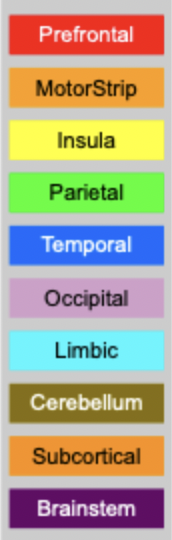

**Supplementary Figure 7.**

NJ-CPM Connectogram. Negative network, edges from SMN nodes, degree threshold = 5.

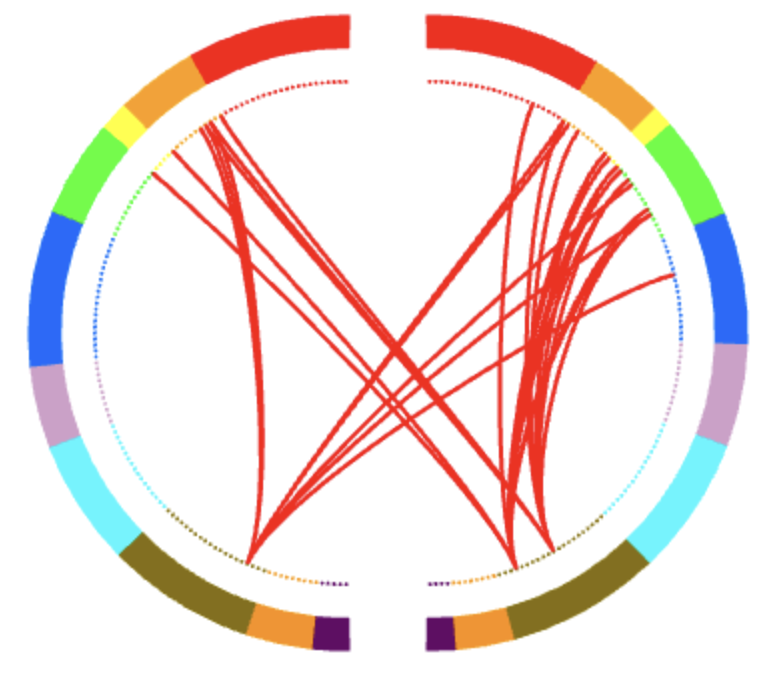

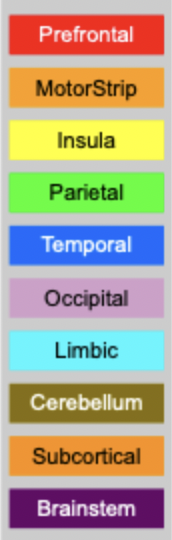

**Supplementary Figure 8.**

AA-CPM Connectogram. Positive network, edges from Unknown nodes, degree threshold = 5.

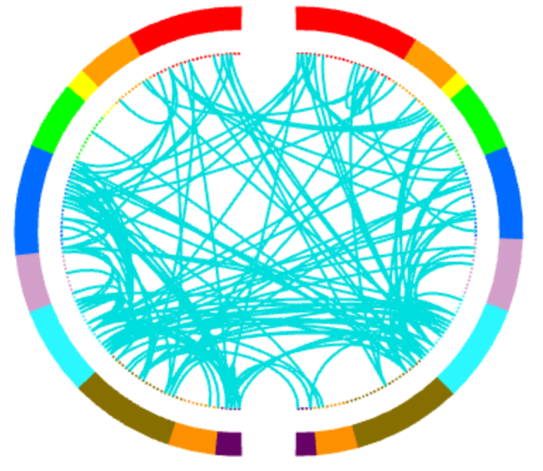

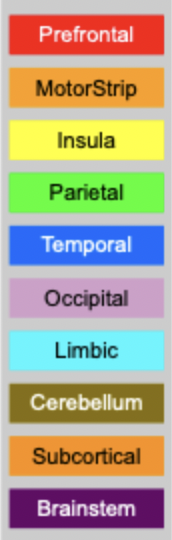

**Supplementary Figure 9.**

AA-CPM Connectogram. Negative network, edges from Unknown nodes, degree threshold = 5.

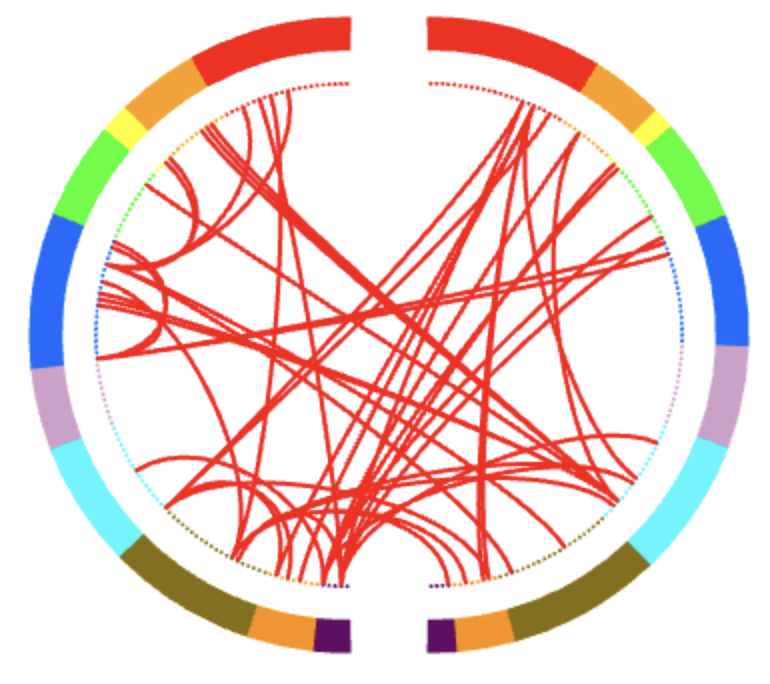

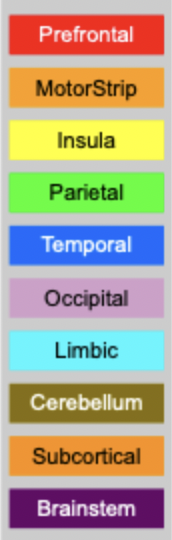

**Supplementary Figure 10.**

NJ-CPM Connectogram. Positive network, edges from Unknown nodes, degree threshold = 5.

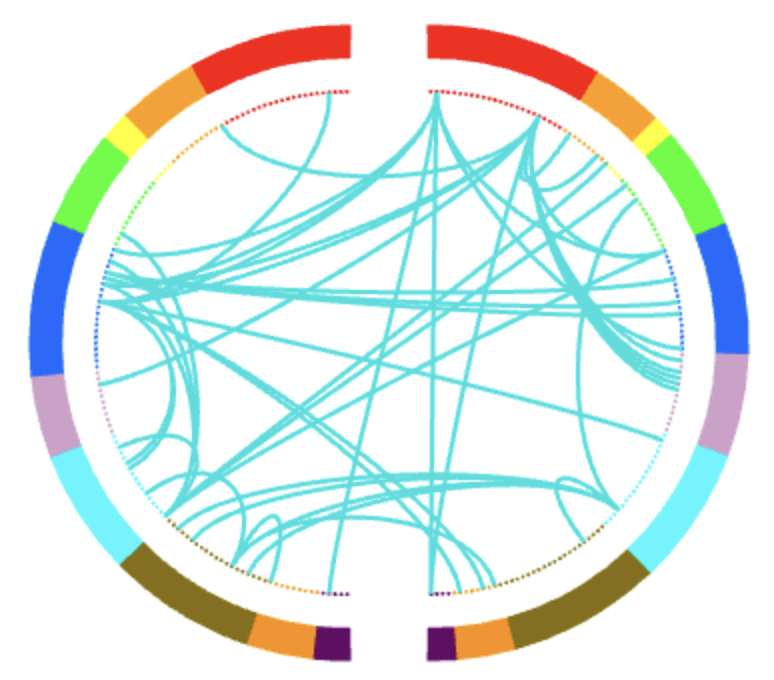

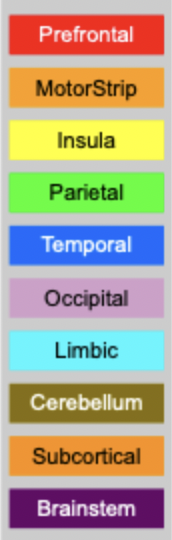

**Supplementary Figure 11.**

NJ-CPM Connectogram. Negative network, edges from Unknown nodes, degree threshold = 10.

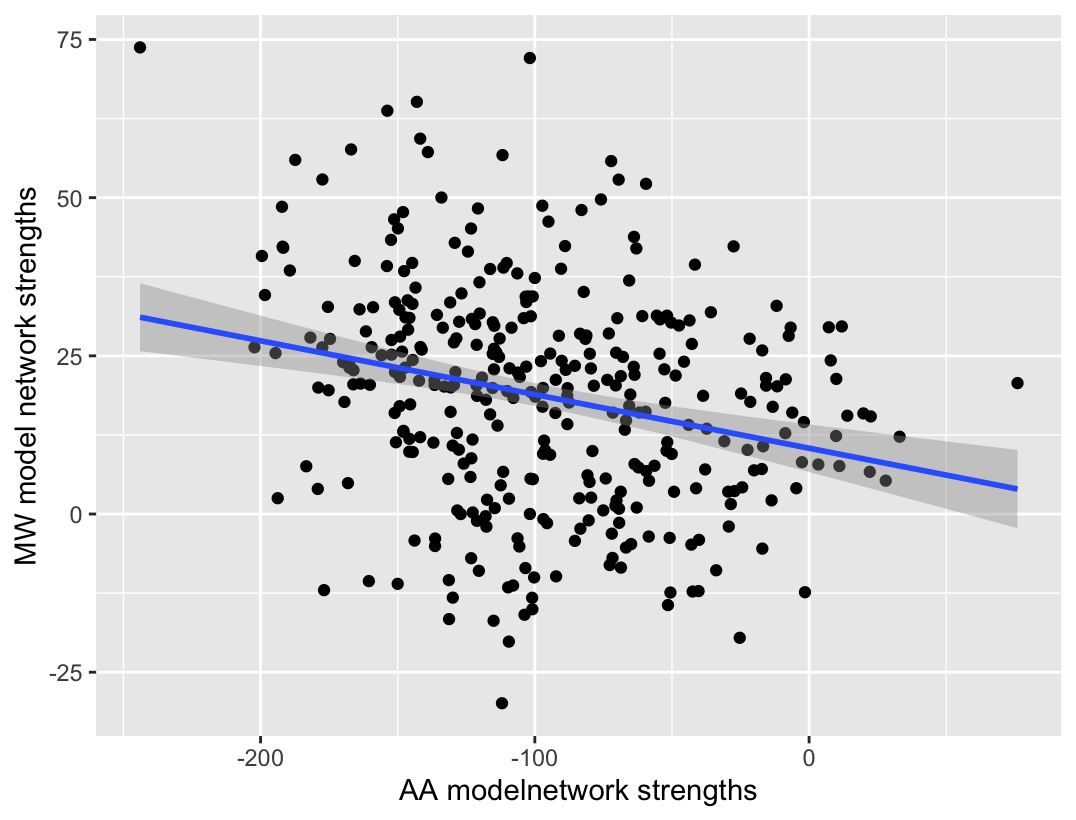

**Supplementary Figure 12.**

Correlations between *Acting with Awareness* model network strengths and literature Mind-wandering model network strengths in all data. Blue line is linear best fit, with 95% gray confidence intervals.

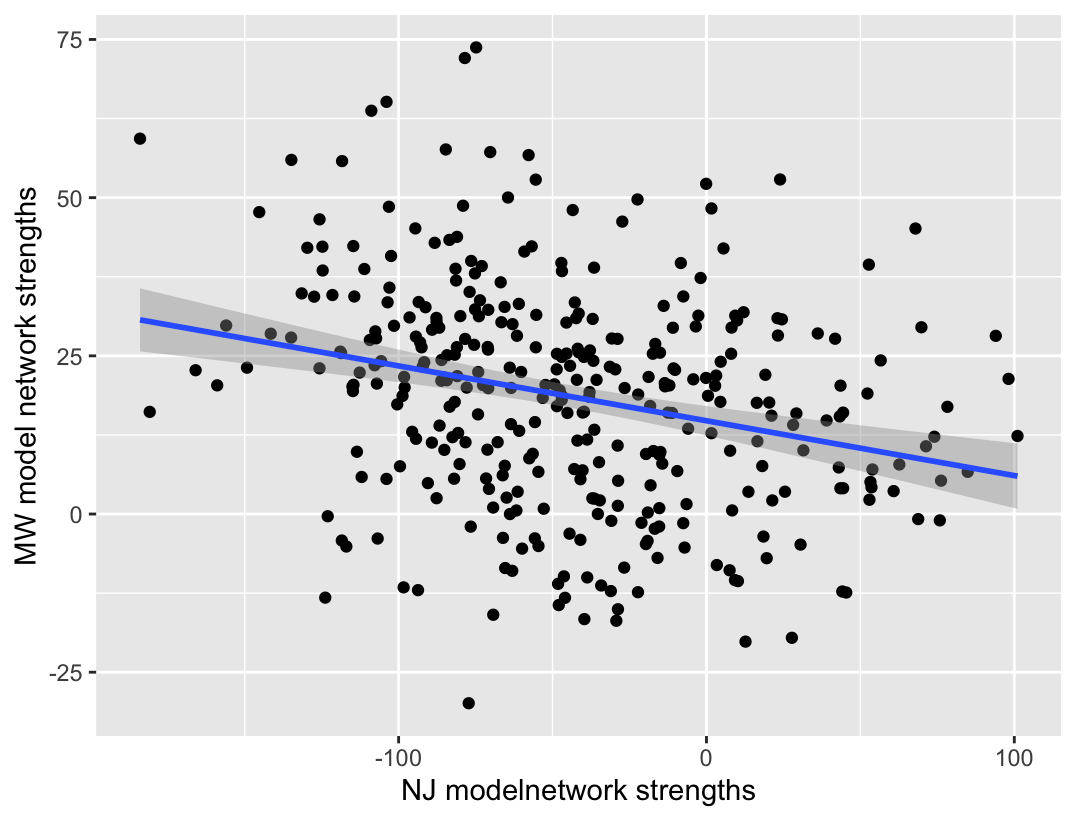

**Supplementary Figure 13.**

Correlations between *Non-judging* model network strengths and literature Mind-wandering model network strengths in all data. Blue line is linear best fit, with 95% gray confidence intervals.

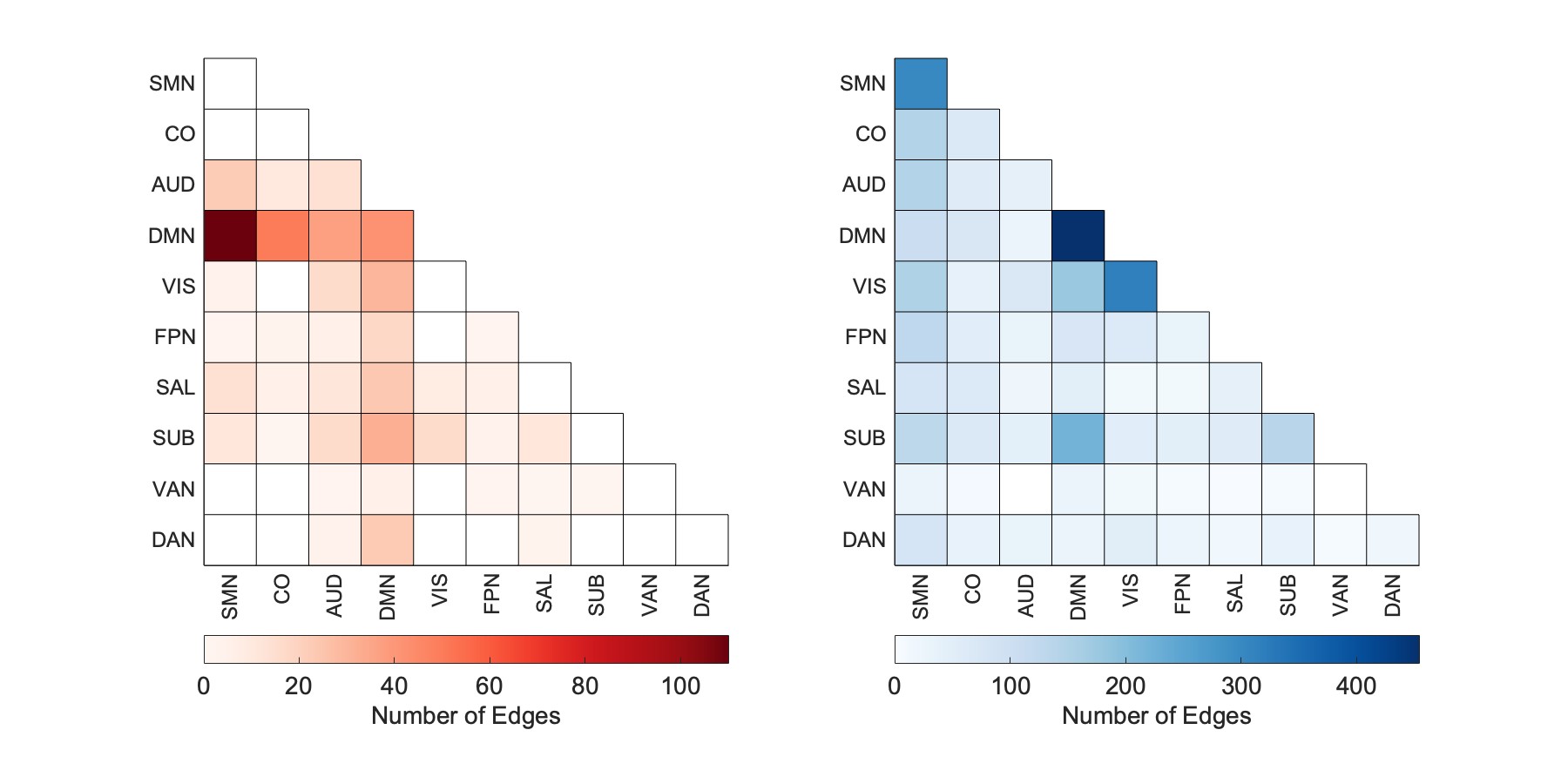

**Supplementary Figure 14.**

NJ-CPM positive network, when trained on shuffled data across sites.

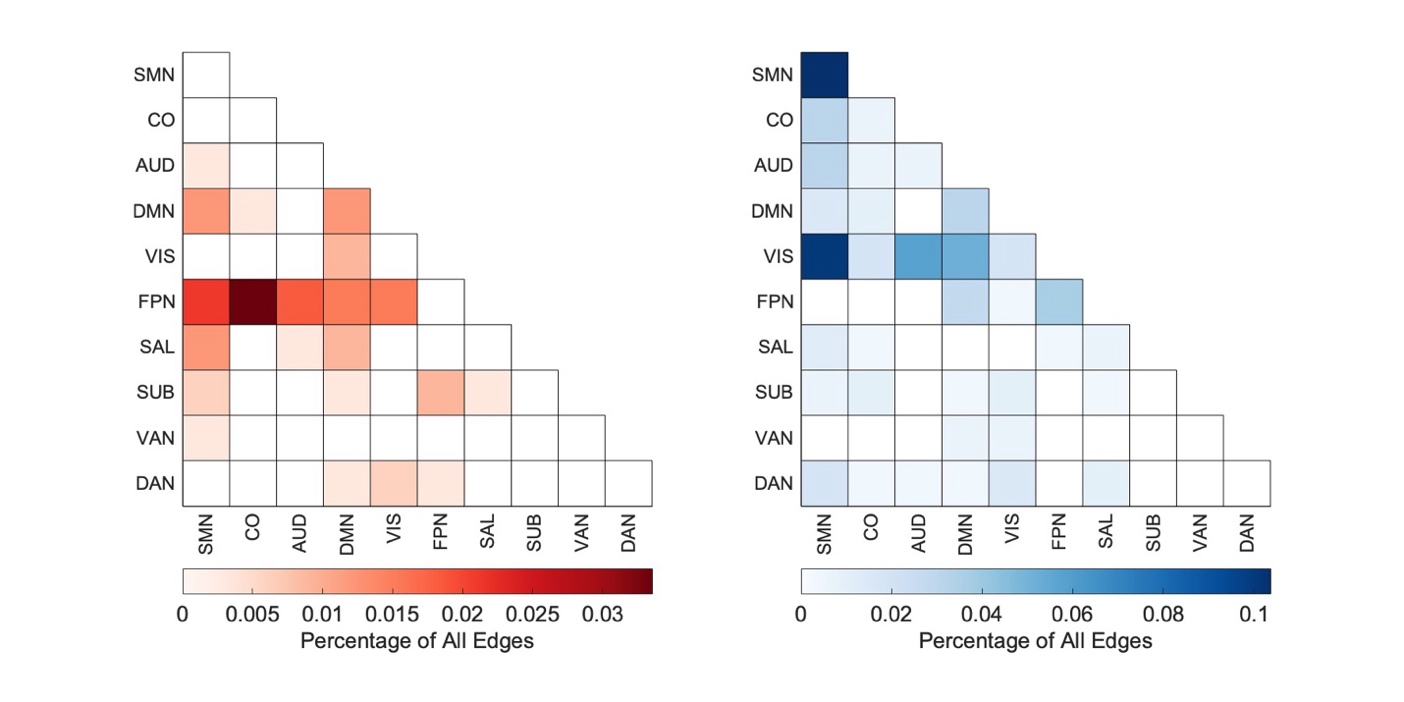

**Supplementary Figure 15.**

Acting with Awareness models, edges normalized by total number of edges. Positive network, in red. Negative network, in blue.

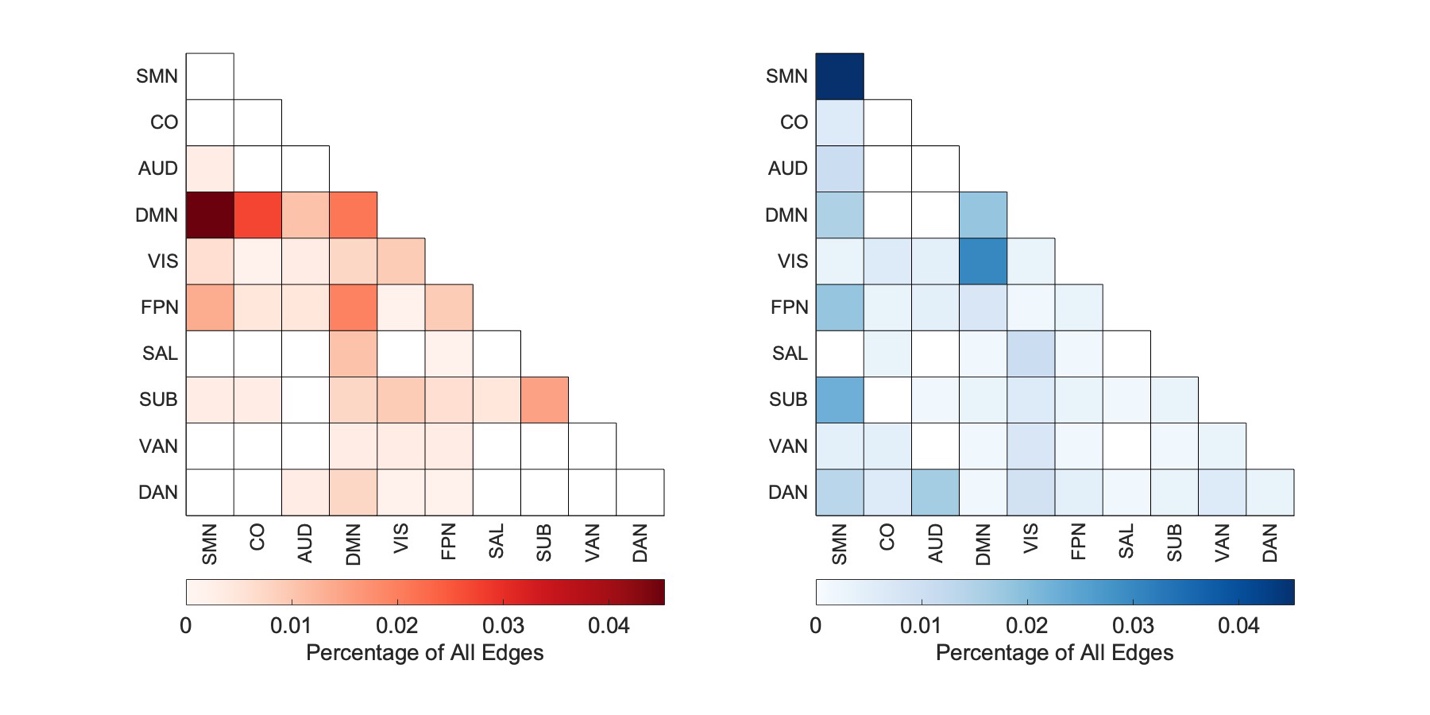

**Supplementary Figure 16.**

Non-judging models, edges normalized by total number of edges. Positive network, in red. Negative network, in blue.

| **Name** | **AA** | **D** | **O** | **NJ** | **NR** | **Total** |
| --- | --- | --- | --- | --- | --- | --- |
| AA |  |  |  |  |  |  |
| D | 0.33**** |  |  |  |  |  |
| O | 0.19** | 0.47**** |  |  |  |  |
| NJ | 0.36**** | 0.22** | 0.14* |  |  |  |
| NR | 0.33**** | 0.34**** | 0.42**** | 0.37**** |  |  |
| Total | 0.66**** | 0.72**** | 0.64**** | 0.64**** | 0.69**** |  |
| TotalNot Observe | 0.72**** | 0.69**** | 0.42**** | 0.71**** | 0.68**** | 0.97**** |

**Supplementary Table 1.**

Correlations between facets of mindfulness in the Wisconsin training dataset. *, *p* < 0.05, **, *p <* 0.01, ****, p* < 0.001, ******, *p <* 0.0001. AA: Acting with Awareness, D: Describing, O: Observing, NJ: Non-judging, NR: Non-reactivity, Total: Total scores, TotalNotObserve: Total without Observing facet.

| **Subscale** | **R** | **p** |
| --- | --- | --- |
| Total | 0.1 | 0.16 |
| TotalNotObserve | 0.15 | 0.1 |
| AA | 0.22 | 0.017* |
| D | 0.17 | 0.074 |
| NJ | 0.21 | 0.025* |
| NR | 0.0015 | 0.49 |
| O | -0.063 | 0.65 |

**Supplementary Table 2.**

Training dataset performance for FFMQ subscales and total scores, permutation tested. R: correlation coefficient, p: p-value. Total: Total scores, TotalNotObserve: Total without Observing facet, AA: Acting with Awareness, D: Describing, NJ: Non-judging, NR: Non-reactivity, O: Observing. *, below selection threshold, p < 0.05.

| **Model** | **Subscale** | **Correlation (parametric)** |
| --- | --- | --- |
| CPM-Tangent | AA | 0.27 |
| CPM-Tangent (negative) | NJ | 0.26 |
| CPM-Tangent | NR | 0.26 |
| CPM-Tangent | Total | 0.26 |
| CPM-Tangent (negative) | D | 0.2 |
| BBS (dim=150) | AA | 0.19 |
| Enet (λ=1, α= 0.1) | NJ | 0.27 |
| Enet (λ=1, α= 1) | TotalNotObserve | 0.25 |
| Enet (λ=0.422, α= 0.775) | D | 0.22 |
| Enet (λ=0.032, α= 0.1) | AA | 0.15 |

**Supplementary Table 3.**

Other predictive modelling approaches. Significant prediction results in the Wisconsin training set are shown, sorted by method and by magnitude of the correlation. CPM: connectome predictive modelling. BBS: brain basis set. Enet: Elastic Net Regression. Tangent: tangent parameterization of covariance matrix. (negative) refers to the negative network of CPM. Dim: dimensionality reduction for BBS. Alpha and lambda are parameters of elastic net (ENet).
